# Cellular and molecular coding of habit-driven behavior in the dorsal raphe nucleus

**DOI:** 10.64898/2026.10.01.755903

**Authors:** Laura Chalençon, Rachel M. Rahn, Revathy U. Chottekalapanda, Alexandra Lenz, Tuo Zhang, Alexander Donatelle, Joseph M. Stujenske, Robert N. Fetcho, Puja K. Parekh, Jacob Roshgadol, Matthew A. Wright, Conor Liston

**Author notes:** These authors contributed equally to this work (Laura Chalençon, Rachel M. Rahn).

## Abstract

Chronic stress has potent effects on reward-seeking behavior, reducing hedonic drive but also biasing individuals toward habitual responding at the expense of goal-directed action. While prior studies have implicated corticostriatal circuits in these shifts, the contribution of the dorsal raphe nucleus (DRN)—a central neuromodulatory hub that regulates motivational states—remains poorly understood. Here, we identify a key role for DRN neurons in mediating stress-induced transitions from goal-directed to habit-driven behavior. Using two-photon calcium imaging in awake behaving mice, we show that DRN neurons encode representations of both effort expenditures and reward acquisition that are integrated at the single-neuron level. Chronic stress selectively enhances reward-related responses in habit-driven individuals, and deficits in reward-related activity in DRN neurons predict future habit formation after chronic stress. Single-nucleus transcriptional profiling revealed that chronic stress drives habit-related gene expression changes in GABAergic DRN neurons, including upregulation of genes implicated in synapse function and inhibitory signaling, and identified potential pharmacological treatment targets. Optogenetic inhibition of DRN GABA neurons rescued stress effects on habit formation, driving stable shifts in operant responding that persisted for days. Together, these findings establish the DRN as a critical modulator of stress-induced behavioral inflexibility, and identify a new role for DRN GABAergic neurons in determining stress susceptibility and driving stable and persistent switches in operant responding during recovery from chronic stress.

---

Stress is a powerful modulator of behavior, exerting widespread effects on motivation, reward processing, and effort valuation^1^. Chronic stress has been shown to disrupt hedonic function, dampening responses to rewarding stimuli while simultaneously enhancing habit-driven behaviors at the expense of goal-directed actions^2,3^. These effects may contribute to the pathophysiology of addictions, depression, obsessive compulsive disorder and other neuropsychiatric conditions, in which stress can exacerbate habit-driven behaviors and undermine adaptive decision-making^4^. Despite decades of research, the precise neural circuits that mediate stress-induced shifts in reward-seeking and habitual behavior remain incompletely understood. Identifying these circuits is critical not only for understanding the neurobiology of stress-related psychiatric disorders but also for delineating fundamental mechanisms by which stress shapes behavior.

Extensive evidence supports the existence of two parallel systems that govern reward-seeking behavior: a goal-directed system, which flexibly updates behavior based on outcome value and contingencies, and a habit-driven system, which promotes stereotyped, automatic responding after repeated conditioning^5–8^. Chronic stress has been shown to alter both systems in distinct ways^2,9^. Stress-induced deficits in hedonic behavior and effort valuation reduce goal-directed responding^10,11^, while simultaneously biasing behavior toward habitual and compulsive responding—a shift that may underlie stress-induced vulnerability to addictions and depression^12,13^. While previous work has largely focused on regions such as the prefrontal cortex, ventral tegmental area, nucleus accumbens, anterior cingulate cortex, and basal ganglia in mediating these effects^1,4^, one region that has been comparatively neglected despite its potential to orchestrate stress-induced changes in reward-seeking and compulsive behaviors is the midbrain dorsal raphe nucleus (DRN).

The DRN, the principal source of serotonin (5-HT) projections to the forebrain, is highly stress-sensitive and plays a central role in modulating stress responses^14,15^. Like other brainstem neuromodulatory centers, it is well positioned to exert powerful effects on a variety of stress sensitive behaviors through widespread projections to distributed cortical and subcortical brain regions^53–56^. Beyond its well-established role in mood regulation, the DRN is strongly implicated in stress-related behavioral adaptations, including shifts in reward valuation and action selection^16^. The DRN itself is exquisitely stress-sensitive, undergoing neurochemical and molecular changes that may underlie its capacity to modulate stress responses^17^. Notably, chronic stress alters DRN neurotransmission, including stress-induced plasticity in corticotropin-releasing hormone receptor expression and neurotransmitter switching^15,18^.

Beyond its role in responding to stress, the DRN is also thought to play a central role in reward-seeking and impulsivity^19,20^. DRN serotonin neurons project to key reward-processing regions, including the nucleus accumbens, prefrontal cortex, and ventral tegmental area, modulating motivation and reinforcement learning^21–23^. The DRN is thus centrally positioned to regulate stress-induced shifts in reward-seeking and habit-driven behaviors. However, despite its critical role in both stress and reward processing, the contribution of the DRN to shaping stress-induced transitions between goal-directed action and habitual behavior remains largely unknown.

To address this gap, we interrogated the role of the DRN in mediating stress-induced changes in reward-seeking behavior. We utilized a progressive ratio task with outcome devaluation, a classic paradigm that is particularly well-suited for this purpose, as it can be performed by head-fixed animals in a physiology-compatible platform and it enables the dissociation of effortful, goal-directed reward-seeking from habitual responding^2,5^. Prior studies have shown that chronic stress reduces effortful goal-directed behavior but also promotes compulsive, habit-driven responses in this task^2,24^. We employed a multifaceted approach to dissect the DRN’s role in these processes. Using two-photon calcium imaging to record single-cell activity in the DRN before and after chronic stress, we examined how stress alters the neural encoding of effort and reward, and how these effects differ in animals that develop compulsive, habit-driven licking compared to those that do not. Next, we used single-nucleus RNA sequencing to identify cell type-specific molecular adaptations in the DRN that may contribute to these behavioral shifts, revealing stress-induced transcriptional changes in GABAergic populations. Finally, we used optogenetic tools to manipulate activity in specific DRN neuronal subtypes and evaluate their necessity for stress-induced habit-driven behavior and their ability to rescue stress effects on habit formation. Together, our results identify a central role for DRN GABAergic neurons in determining stress susceptibility, enhancing habit formation in chronic stress states, and driving stable and persistent switches in operant responding during recovery from chronic stress.

## RESULTS

### Dorsal Raphe Neurons Integrate Effort and Reward Signals

Effort valuation is a process for integrating information about anticipated rewards and the effort required to obtain them, and regulating decisions to expend effort to obtain a reward^25,26^. Previous work indicates that DRN neurons are sensitive to rewards and emotionally salient experiences, and that they regulate reward-seeking behavior and reinforcement learning^27–30^. However, whether and how DRN neurons encode effort requirements and integrate this information with representations of rewarding experiences is not well understood. To address this question, we used two-photon calcium imaging and chronically implanted GRIN lenses to record the activity of populations of DRN neurons with single-cell resolution while mice performed a progressive ratio task—a classic paradigm for evaluating motivation and habit formation in rodents^31^.

To this end, we developed a physiology-compatible platform for implementing a progressive ratio task in head-fixed mice (**Fig. 1a**), which learned to lick a spout repeatedly to obtain water rewards, first during a 5-day fixed-ratio training period (1:1 lick : reward ratio; 136 trials per 30-minute session; **Fig. 1b**). This was followed by 5 days of progressive ratio testing, in which the number of licks required to obtain the reward increased exponentially across trials (from 1:1 to 250:1 across up to 80 trials in 30 minutes). Motivation to work for reinforcement was assessed in terms of the breakpoint, or maximum number of licks performed during a trial before the subject stops responding. In contrast to the fixed ratio sessions, in which most mice achieved the maximum of 136 rewards within 30 minutes, all mice stopped responding prior to the end of the session during the progressive ratio phase (mean breakpoint = 91 ± 52 licks over 54 ± 15 trials completed; **Fig. 1b**). To further validate the task, we confirmed that lick frequency during the effort expenditure phase of the task, but not during the reward acquisition phase, was negatively correlated with effort requirements (**Fig. 1c**).

**Figure 1:**
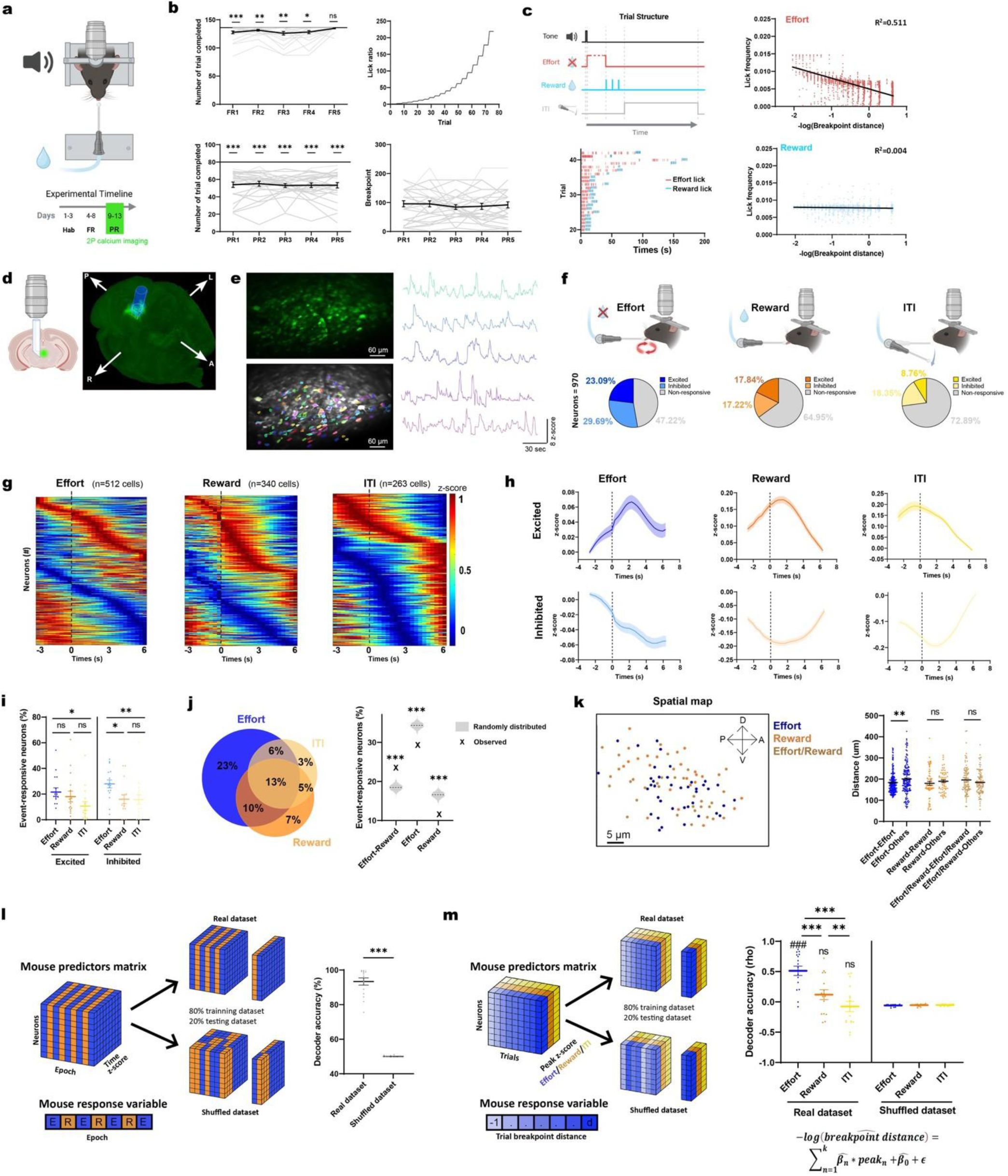
Integrated encoding of effort and reward signals in dorsal raphe neurons. **A.** Task schematic and experimental timeline. Hab: habituation, FR: fixed ratio, PR: progressive ratio. **B.** During the fixed ratio period (FR, **upper left**), mice learned to increase their licking over time to obtain more rewards (n=25 mice, p<0.001 by Friedman test for repeated measures) and consistently reached the maximum number of achievable trials by session 5 (asterisks denote significant difference from maximum achievable trials by one-sample Wilcoxon). During the progressive ratio period (PR), licking requirements increased with repeated trials (**upper right**), licking behavior did not increase over time (**lower left**, n=25 mice, p=0.989 by Friedman test), and mice did not reach the maximum number of achievable trials, disengaging from the task at a breakpoint of ∼100 licks (**lower right,** equivalent to ∼60 trials). **C.** Trial structure (**upper left**) and representative lick timestamps (**lower left**). Lick frequency during the effort period (**upper right**), but not during the reward acquisition period (**lower right**), was anticorrelated with the breakpoint distance (linear regression, n=3,873 trials, R^2^=0.511, p<0.001). **D.** 2P calcium imaging schematic. A: anterior, P: posterior, R: right, L: left. **E.** Representative field of view (mean projection over time), cell segmentation, and calcium traces. **F.** Proportion of neurons excited (significantly positive beta weight in GLM) or inhibited (negative weight) by effort, reward or ITI over N=970 imaged neurons in 16 mice. **G.** Trial-averaged activity histograms for DRN cells. Each cell’s mean trace is normalized to its peak value, and cells are sorted by the time of peak excitation (top) or inhibition (bottom). **H.** Mean activity traces (+/- SEM) time-locked to effort, reward, or ITI onset, and averaged across all trials and all neurons with significant beta weights in (G). **I.** The proportion of neurons with activity modulated by effort was significantly higher than those with activity modulated by reward consumption or ITI (n=16 mice, linear mixed-effects model: event effect F(2,30)=8.916, P<0.001; * = P<0.05, ** = P<0.01, Bonferroni corrected). **J.** Distribution of neurons responsive to one or more task features. The percentage of multimodal neurons coding for both effort and reward is significantly higher than expected by chance, and the proportions of unimodal neurons are significantly lower than expected, given the observed proportion of cells encoding each event (p<0.001 in simulations of null distribution, see Methods). **K.** Representative map showing spatial distribution of effort-, reward-, and effort/reward-coding cells (**left**). The mean distance between a pair of effort-coding neurons was lower than the distance between an effort-coding neuron and a neuron coding for another event (**right**, n=225 effort neuron pairs, n=65 reward neuron pairs, n=101 multimodal effort/reward coding neurons, linear mixed-effects model, Bonferroni corrected for post-hoc comparisons: ** = p < 0.003, NS = not significant. **L.** Schematic for decoding task epoch from DRN neuronal activity (**left,** E: effort, R: reward). Decoder accuracy (n=15 mice) was higher than that observed in shuffled data (**right,** Wilcoxon, P<0.001). **M.** Schematic for predicting breakpoint distance from DRN neuronal activity during the effort, reward, or ITI period (**left**). Decoder accuracy was higher for models trained on DRN activity during the effort period than the reward or ITI period, and significantly higher than expected by chance in shuffled data (mixed-effects model: effect of task epoch, F(2,30)=12.49 p<0.001; real vs. shuffled data, F(1,15)=16.85 p=0.003). # = significantly different from shuffled data, p<0.001; NS = not significantly different from shuffled data. *** = significantly different by task epoch, p<0.003. Swarm plots represent mean (line) ±SEM and individual data points (dots). Line plots represent mean (line) ±SEM (shaded area). Scatter plots represent individual data points (dots). Violin plots represent mean (dashed lines), first and third quartiles (dotted lines). (n=16 mice)

Next, we recorded the activity of DRN neuronal populations during progressive ratio task performance, using a viral vector to express a genetically encoded calcium indicator (AAV-hSyn-GCaMP6s) in DRN neurons. To facilitate two-photon calcium imaging of large neuronal populations with minimal damage to the DRN, we implanted a customized microendoscope consisting of a GRIN lens ending in a prism facing laterally into the DRN, which enabled the simultaneous imaging of up to 150 neurons per mouse along the entire anteroposterior and dorsoventral axes of the DRN (**Fig. 1d; Supplementary Fig. 1a-e**). We also adapted existing preprocessing pipelines for two-photon calcium imaging to overcome technical challenges specific to imaging the DRN, including customizations for motion correction and denoising using DeepCadRT^32^ (see STAR Methods).

To characterize the coding properties of individual neurons, we used a generalized linear model (GLM) to identify neurons responsive to the effort expenditure, reward acquisition, and intertrial interval (ITI) periods of the task (**Fig. 1c; Supplementary Fig. 1f**). Activity in 88.8% of DRN neurons (861 out of 970 cells recorded) was significantly modulated by at least one task feature (**Fig. 1f**). Interestingly, most DRN neurons responded preferentially to just one task feature. As in previous work^30^, ∼35% of DRN neurons were responsive to reward acquisition, with roughly equal numbers of cells activated or inhibited by reward, but an even larger population of neurons (∼53% of all recorded cells) were responsive to effort (**Fig. 1f-i**). Together, these data show that DRN neuronal populations are sensitive not only to reward consumption but also to effort expenditures.

Effort valuation computations are understood to require integrated representations of both reward- and effort-related signals. To better understand how effort- and reward-related information is represented in the DRN, we tested whether these signals are encoded by distinct neuronal populations or are computed within the same neurons. We found that activity in ∼23% of recorded neurons was significantly modulated by both effort expenditures and reward consumption—a proportion significantly higher than expected by chance based on the observed proportion of cells encoding each event (**Fig. 1j**). Based on prior work identifying anatomically defined subpopulations of DRN neurons that encode reward- or anxiety-related stimuli with distinct projection targets^30^, we also tested whether effort- and reward-responsive neurons were topographically segregated. Although effort- and reward-coding neurons did not clearly differentiate along the dorsoventral or anteroposterior axes (**Supplementary Fig. 1g**), we found that effort-coding neurons were spatially clustered, while the others were not (**Fig. 1k**). These results show that effort-related information is topographically organized within DRN, and that effort and reward signals are integrated within DRN neurons at the single-cell level.

Finally, we asked whether DRN neuronal ensembles encode sufficient information at the population level to decode key task features and decisions. To test this, we trained support vector machine (SVM) classifiers to decode whether the mouse was engaged in effort expenditure or reward acquisition at any given moment during the task, based on the population-level activity of DRN neurons (for details, see STAR Methods). SVM decoders successfully differentiated between the effort and reward acquisition task epochs with ∼90% accuracy (vs. ∼50% or chance prediction in shuffled data), indicating a highly robust population-level representation of both task epochs (**Fig. 1L**). This, in turn, motivated us to ask whether DRN cells also encode a population-level signal that predicts future decisions to discontinue expending effort to obtain a reward. To test this, we trained decoders to predict the future breakpoint in a given session (**Fig. 1m**). In accord with our hypothesis, peak activity during the effort expenditure period—but not during the reward acquisition period—predicted the future breakpoint (**Fig. 1m**). Together, these data show that DRN neurons integrate effort- and reward-related signals at the single-cell level, and encode future decisions to disengage from the task.

### Reward activity in DRN neurons predicts future habit formation after chronic stress

Previous work has shown that chronic stress disrupts numerous features of reward-seeking behavior and reduces willingness to expend effort to obtain rewards^10,11^. However, other studies indicate that stress may *increase* habit-driven operant responding, especially under experimental conditions involving extensive task experience, which are more likely to give rise to habits^2,9^. How stress affects DRN function in this context is unknown. To investigate how chronic stress influences DRN function, we used two-photon calcium imaging to record the activity of DRN neurons during progressive ratio task performance, assessed before and after an extensively validated, 10-day chronic social defeat stress paradigm (**Fig. 2a**). Chronic stress caused a significant increase in the breakpoint (**Fig. 2b**), as well as expected changes in grooming, social interaction, and avoidance behavior that we used to validate the chronic stress model (**Supplementary Fig. 2a-c**). We also observed individual differences in changes in operant responding after chronic stress: increased responding at the group level was driven by a stress susceptible subset animals (∼57%, 8 of 14 mice) that showed a significant increase in operant responding as measured by increased breakpoint at the individual level, whereas the remaining 43% showed no significant change in breakpoint.

**Figure 2.**
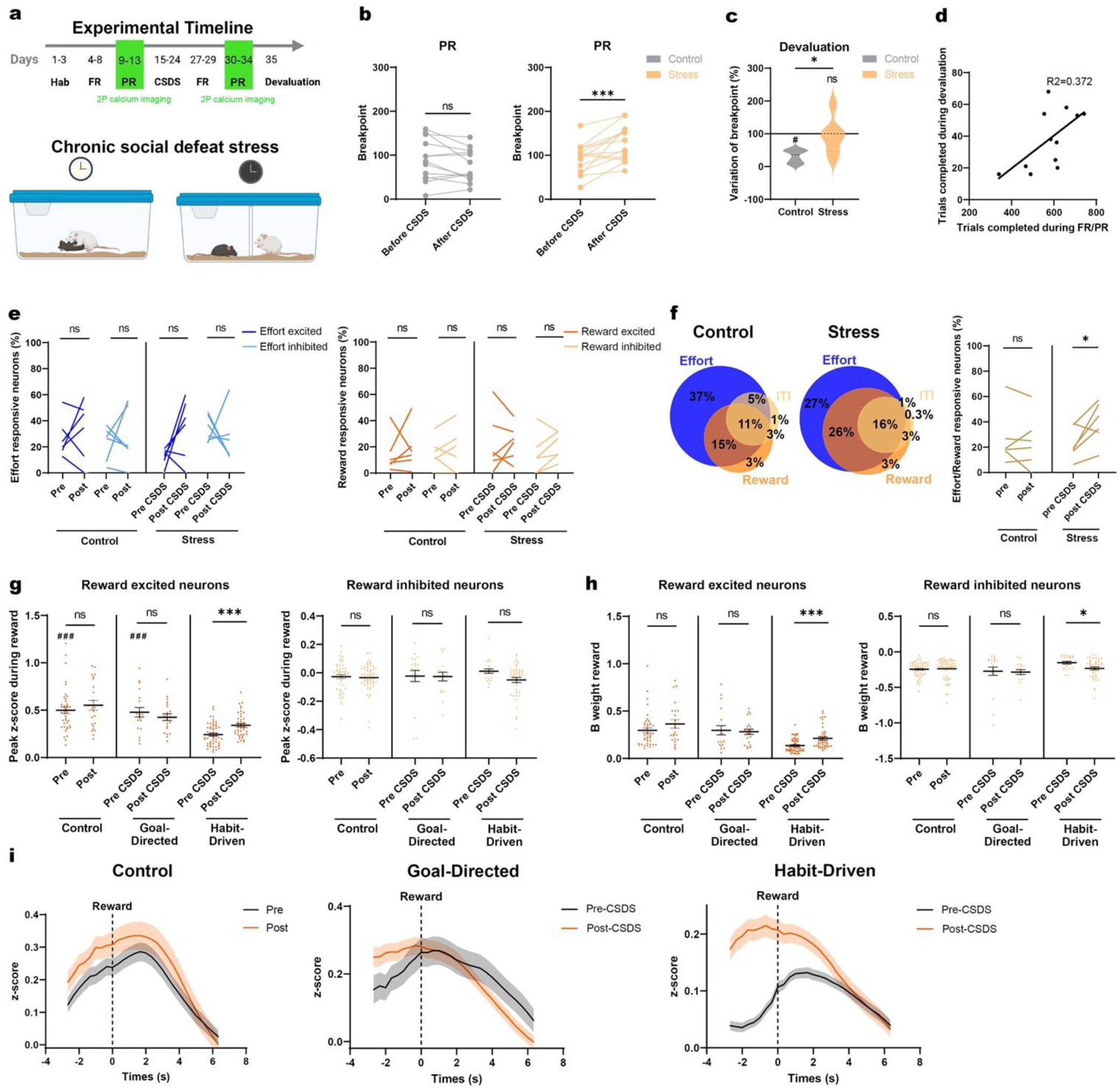
Reward activity in DRN neurons predicts future habit formation after chronic stress. **A.** Timeline for chronic social defeat experiment. Hab: habituation, FR: fixed ratio, PR: progressive ratio, CSDS: chronic social defeat stress. **B.** Social defeat stress (CSDS) increased the breakpoint in stressed mice (**right,** n=14 mice, Wilcoxon, p<0.001), which was stable over time in unstressed control mice (**left,** n=13 mice, Wilcoxon, p=0.076). **C.** Reward devaluation decreased the breakpoint in control but not in stressed mice, an indicator of habit-driven responding (n=6/group; Mann-Whitney: Control vs Stress, p=0.013; one-sample Wilcoxon testing difference from 100% within each group: control, p=0.031, Stress, p=0.999). Violin plots in represent mean (dashed lines), first and third quartiles (dotted lines). **D.** The total number of trials completed after CSDS during FR and PR sessions was correlated with the number of trials completed during the devaluation session (n=12 mice, R^2^=0.372, p=0.035). **E.** CSDS had no effect on the proportion of neurons coding for effort or reward (linear mixed effects model, treating subject as a random effect; n=6 mice/condition; main effect of group: F(1,20)=0.018, p=0.895; main effect of stress: F(1,20)=1.640 p=0.208; group x stress interaction: F(1,20)=2.055, p=0.160. NS = not significant). **F.** Distribution of task feature-responsive neurons by stress condition (**left)**. The percentage of multimodal neurons coding for effort and reward increased after stress but not in unstressed controls (**right**; linear mixed effects model, n=6 mice/condition; group x stress interaction: F(1,10)=5.923 p=0.035; pre-vs. post-CSDS post-hoc contrast: p=0.021, Bonferroni corrected. NS = not significant). **G.** Effect of CSDS on peak activity in reward-coding neurons. In stress-susceptible, habit-driven mice, peak activity during reward acquisition increased after CSDS in reward-excited neurons (n=55 cells pre-CSDS and n=23 cells post-CSDS in 6 mice; ***P<0.001, Mann-Whitney, Bonferroni corrected). Prior to CSDS, peak activity during reward acquisition was also lower in these mice compared to unstressed controls (pre n=42 cells and post n=23 cells in 6 mice) and stress-resilient, goal-directed mice (pre-CSDS n=19 cells and post-CSDS n=21 cells in 3 mice; ### P<0.001, Mann-Whitney, Bonferroni corrected). There were no other significant effects of stress on reward-related activity (NS). Swarm plots represent mean (line) ±SEM and individual data points (dots). **H.** Analysis of the reward-associated GLM beta weight confirmed the pattern observed in (G): increased reward-related activity after CSDS in reward-excited cells in stress-susceptible, habit-driven mice (n=55 cells pre-CSDS and n=23 cells post-CSDS in 6 mice; ***P<0.001, Mann-Whitney, Bonferroni corrected), as well as a modest decrease in reward-related activity in reward-inhibited cells in this group (*P=0.039, Mann-Whitney, Bonferroni corrected). **I.** Mean activity traces (+/- SEM) time-locked to reward onset, and averaged across all trials and all neurons from (G-H) with significantly positive reward-associated beta weights in unstressed controls (**left**), stress-resilient / goal-directed mice (**middle**), and stress-susceptible / habit-driven mice (**right**).

To differentiate stress effects on goal-directed reward seeking versus habit-driven operant responding, we devalued the reward by providing water access immediately prior to a final test session. Unstressed control mice adapted their behavior to reward devaluation with a significant reduction in the breakpoint, but chronically stressed mice did not (**Fig. 2c**), indicating a failure to adapt to changing reward values at the group level and an increased propensity for habit formation. They also suggest that the increased breakpoint observed after chronic stress was explained in part by increased habit-driven licking, as opposed to goal-directed reward seeking. Indeed, breakpoint behavior on the reward devaluation day was significantly correlated with the total number of trials completed during the fixed and progressive ratio phases (R^2^ = 0.372; **Fig. 2d**), lending further support to the interpretation that extensive task exposure drove habit formation in chronically stressed, but not unstressed control, mice. Accordingly, individual differences in breakpoint after chronic stress were strongly predictive of a subsequent failure to adapt to reward devaluation (χ^2^=4.38, P=0.036), such that 86% of stress-susceptible mice with increased breakpoint behavior after chronic stress also exhibited habit-driven responding in the reward devaluation test, indicating that individual differences in stress susceptibility in Fig. 2b are stable and predictive of subsequent habit-driven responding.

The circuit-level mechanisms underlying increased habit-driven behavior in chronic stress states are not well understood. To investigate them, we first tested whether chronic stress altered the neuronal responses of DRN neurons to progressive ratio task features. We found that chronic stress had no effect on the proportion of neurons with activity that was significantly increased or inhibited by either effort expenditures or reward acquisition (**Fig. 2e**). However, the proportion of DRN neurons encoding both effort and reward increased after chronic stress, which expanded a multimodal neuronal population responding to both effort and reward (**Fig. 2f**). These results suggest a stress-induced enhancement of the integration of effort and reward signals within the same DRN neurons.

Next, we tested whether stress influenced the degree to which effort- and reward-responsive neurons were modulated by task features and if these effects differed in individuals prone to habit formation after chronic stress. We found that stress increased reward-related activity but only in stress-susceptible, habit-prone mice (**Fig. 2g**).There was no effect of stress on reward-responsiveness in stress-resilient, goal-directed individuals, and there was no effect on effort responsiveness in either group (**Fig. 2h-i; Supplementary Fig. 2d-f**). Importantly, we also found that reward responsiveness prior to stress predicted the subsequent formation of habit-driven responding after stress: specifically, reward-related activity was significantly *reduced* during operant responding prior to stress in mice that went on to develop habit-driven responding, compared to those that did not, and compared to unstressed controls (**Fig. 2g-i).** In summary, these data show that chronic stress increases habit formation, and enhances reward-related responses in DRN neurons in habit-driven individuals but not in individuals with stress-resilient, goal-directed responding. Furthermore, a relative deficit in reward-related responses in DRN neurons prior to stress may predispose individuals to habit formation in response to chronic social stress.

### Chronic Stress Drives Habit-Related Transcriptional Changes in GABAergic DRN neurons

The molecular and cellular mechanisms mediating stress effects on habit-driven behavior—a potential path to identifying treatment targets—are not well understood, particularly in the context of DRN function. Furthermore, DRN neurons are genetically and functionally heterogeneous^33,34^, and it is unknown whether chronic stress engages molecular signaling pathways in specific cell types that predispose to habit formation. To address this, we used single nucleus RNA sequencing to understand how chronic stress influences gene expression in DRN neurons following the same behavioral testing paradigm used above (**Fig. 3a**). DRN tissue was dissected the day after the reward devaluation testing session, and single-nuclei suspensions were prepared for RNA-seq (**Fig. 3a; Supplementary Fig. 3a**).

**Figure 3.**
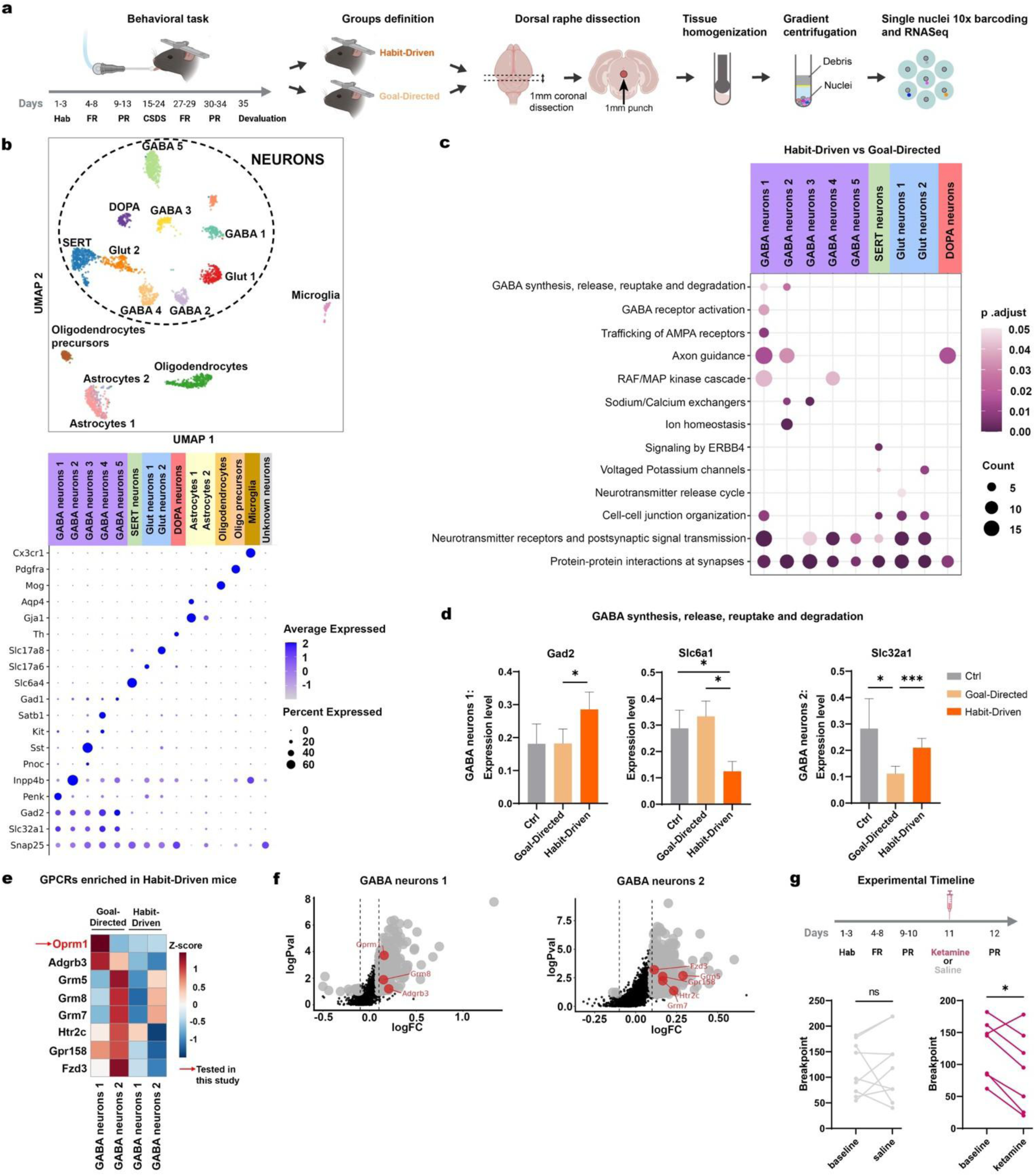
Chronic stress drives habit-related transcriptional changes in DRN GABA neurons. **A.** Schematic of the behavioral task, DRN dissection, and preparation of single-nuclei suspensions for RNA-seq. **B. Upper panel:** UMAP visualization of single-nuclei clusters. **Lower panel:** Dot plot displaying expression of genes used to identify each cell class. The color of each dot indicates the average expression of each gene, and the size indicates the fraction of cells expressing the gene. **C.** Top enriched pathways of differentially expressed genes identified in stress habit-driven vs goal-directed mice. The color of each dot indicates the adjusted p-value of each pathway, and the size indicates the number of significant genes in the pathway. **D.** Bar plots representing the expression level of glutamate decarboxylase 2 (**left**), a key enzyme involved in GABA synthesis; *Slc6a1*, a GABA transporter protein (**middle**); *Slc32a1*, the vesicular GABA transporter (**right**). ***P<0.005, *P<0.05 by Wilcoxon rank sum test. **E.** Heatmap of Z-scores for GPCRs enriched in GABA cell types 1 and 2 in habit-driven mice. **F.** Volcano plots showing differentially expressed GPCRs in habit-driven versus goal-directed mice in GABA cell types 1 (**left**) and 2 (**right**). **G.** Effect of ketamine (30 mg/kg I.P.) on PR breakpoint. **Upper panel:** schematic of experimental timeline. **Lower panel:** PR breakpoint decreases after ketamine (n=7 mice, Wilcoxon rank sum test, p=0.031) but not saline (n=9 mice, Wilcoxon, p=0.570).

After quality control and preprocessing, single nuclei were grouped according to their gene expression profiles using graph-based clustering (see Methods). Unsupervised clustering revealed 15 cell clusters with distinct gene expression signatures. In accord with recently published work, these included serotonergic (SERT) neurons, dopaminergic (DOPA) neurons, two glutamatergic cell types, and five GABAergic cell types, among others (**Fig. 3b**). There were no significant differences in the relative abundance of these cell types across the three experimental groups (**Supplementary Fig. 3b-c**).

Next, we tested for genes that were differentially expressed in stress-susceptible, habit-driven mice, stress-resilient, goal-directed mice, and unstressed control mice (**Supplementary Fig. 4**). Comparing the stress-susceptible, habit-driven mice versus the stress-resilient, goal-directed group, the highest number of significantly altered pathways were found in DRN GABAergic neurons (GABA1 and GABA2 in **Fig. 3c**). Gene set enrichment analysis revealed a host of transcriptional changes involving GABA signaling and synapse function (**Fig. 3c**). Interestingly, a host of differentially expressed genes were also observed in the contrast of the stress-susceptible, habit-driven group vs. unstressed controls, but there were very few differences between stress-resilient, goal-directed mice and unstressed controls (**Supplementary Fig. 5a**).

To further probe how GABA signaling is modulated by chronic stress in habit-driven mice vs goal-directed mice, we examined specific differentially expressed genes with known functions (**Fig. 3d; Supplementary Fig. 5 b-c**). In habit-driven mice, specifically in subsets of DRN GABAergic neurons. chronic stress was associated with increased expression of *Gad2*, which encodes glutamate decarboxylase 2, a key enzyme involved in activity-dependent GABA synthesis; increased expression of *Slc32a1*, the transporter that packages GABA into synaptic vesicles; and decreased expression of *Slc6a1*, which encodes a transporter responsible for the reuptake of GABA from the synaptic cleft. In addition, chronic stress increased the expression of genes encoding GABA receptor subunits (**Supplementary Fig. 5d-e**) Together, these data suggest that habit-driven behavior after chronic stress is associated with a host of transcriptional changes in GABAergic DRN neurons contributing to enhanced GABA signaling and activity.

We also tested for differential expression of genes encoding G-protein coupled receptors (**Fig. 3e,f)**, which are especially appealing drug targets because they are engaged by a large number of existing drug compounds and signal through four common classes of heterotrimeric G proteins with known effects on neuronal activity. Chronic stress increased the expression of at least eight genes encoding GPCRs in DRN GABA neurons in habit-driven mice, including the serotonin 2c receptor and multiple metabotropic glutamate receptors. The most significant effect involved increased expression of *Oprm1* (**Fig. 3f)** which encodes the mu opioid receptor (MOR), a Gi/o-coupled GPCR that reduces neuronal excitability and has been extensively implicated in addictions and habit formation.

In conjunction with data in Fig. 2 revealing increased reward-related activity in DRN cells after chronic stress in susceptible, habit-driven individuals, these results suggest that pharmacological strategies targeting excessive activity in DRN GABAergic neurons, either through MOR signaling or other mechanisms, could be a promising treatment strategy for reducing habit-driven behavior. As a preliminary test of this hypothesis, we asked whether ketamine, a partial MOR agonist that also modulates serotonergic signaling^35^ and may be useful for treating alcohol, opioid, and cocaine addictions^36^, influences operant licking behavior in the progressive ratio task. In accord with our hypothesis, we found that a single subanesthetic dose of ketamine (30 mg/kg, I.P.) caused a significant reduction in the progressive ratio breakpoint in 86% of mice when assessed one day after treatment (**Fig. 3g**).

### Optogenetic Inhibition of DRN GABA Neurons Rescues Stress Effects on Habit Formation

The experiments above revealed how DRN neurons are hyper-responsive to rewards after chronic stress and that this effect is especially pronounced in stress-susceptible, habit-driven individuals. Furthermore, transcriptomic profiling in Fig. 3 showed that differentially expressed genes in habit-driven vs. goal-directed mice tend to localize to GABA cells and involve gene expression pathways that implicate increased GABA cell activity. Thus, we hypothesized that habit-driven behavior in chronically stressed mice may be driven by excessive reward-related activity in DRN GABA neurons and that optogenetic inhibition of GABA cells would rescue stress effects on habit-driven behavior.

To test this, we injected an inducible anion-conducting channelrhodopsin (IC++) in the DRN of Gad2/Cre transgenic mice (or wild type controls) to express this inhibitory opsin specifically in DRN Gad2 neurons and implanted an optical fiber at the same location (**Fig. 4a; Supplementary Fig. 6a,d**). Mice were tested on the same behavioral paradigm as described above, involving progressive ratio testing before and after chronic stress, followed by optogenetic inhibition of Gad2 neurons during progressive ratio task performance (**Fig. 4b**). As predicted, inhibition of DRN Gad2 neurons decreased the PR breakpoint (**Fig. 4c**). To determine whether the inhibition of Gad2 neurons had a long-lasting effect on operant responding, we re-tested mice on three additional PR sessions after the optogenetic testing day but without additional optogenetic stimulation. We found that optogenetic inhibition of DRN Gad2 neurons induced a sustained reduction in operant responding (reduced breakpoint) that persisted for at least 1-2 days (**Fig. 4d**). Remarkably, Gad2 inhibition also fully restored the capacity to adapt to reward devaluation after chronic stress when tested four days later (**Fig. 4e**).

**Fig. 4.**
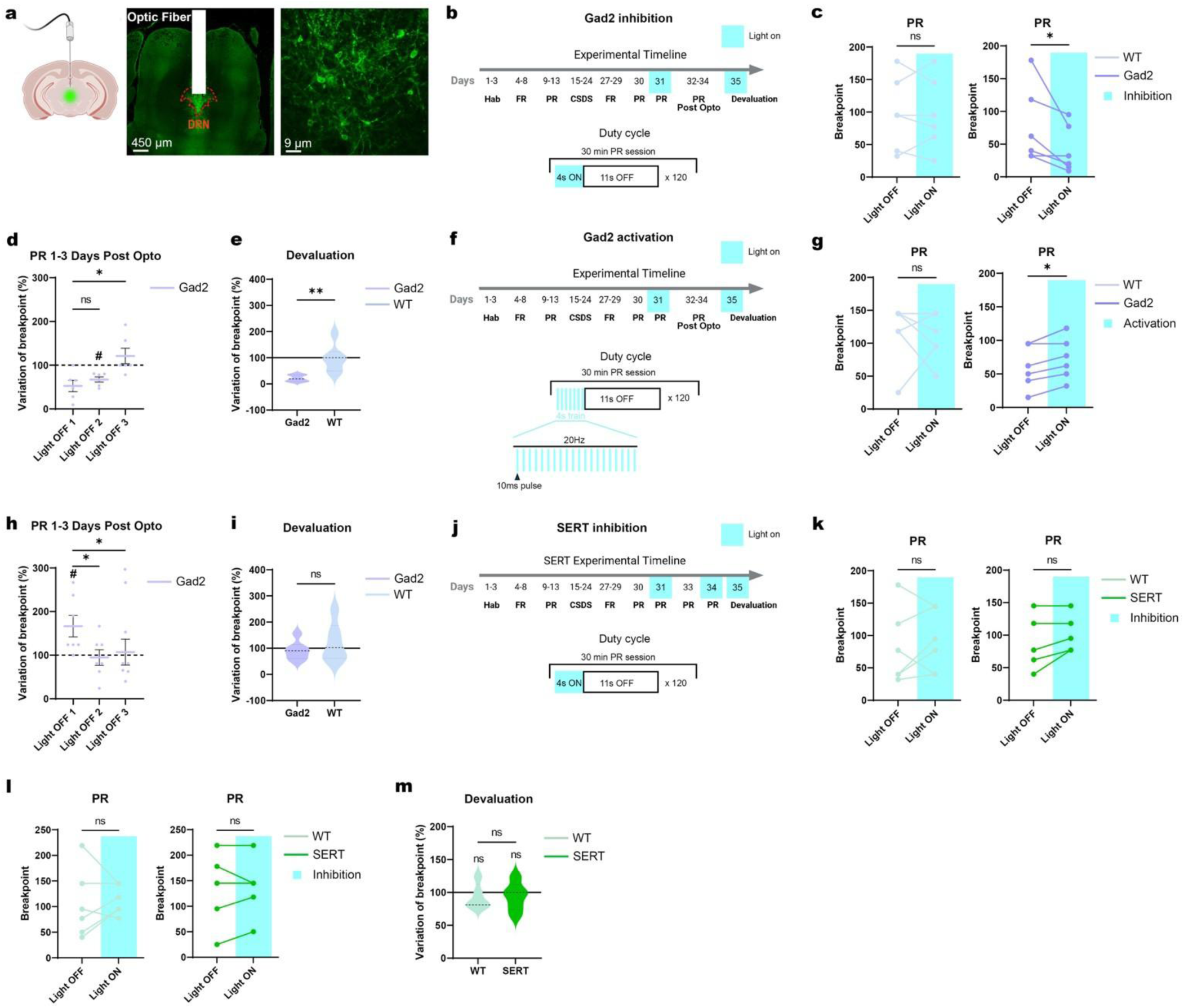
Optogenetic inhibition of DRN GABA neurons rescues stress effects on habit formation. **A.** Schematic (**left**) and representative images of fiber placement (**middle**) and virus localization (**right**). **B.** Experimental timeline for Gad2 cell inhibition using an inducible anion-conducting channel rhodopsin (IC++) in Gad2/Cre mice. **C.** Inhibition of Gad2 neurons decreased the breakpoint in Gad2/Cre mice (N=6 mice/group; *P=0.031, by Wilcoxon rank sum test in Gad2/Cre mice; NS = not significant in wildtype controls). **D.** This effect persisted for 2 days after the optogenetic inhibition session before dissipating on day 3 (p=0.005 by Friedman test for repeated measures; *P=0.012 for Dunn’s post-hoc contrast of Day 3 vs. Day 1; NS = not significant for other post-hoc contrasts. # = significantly different from pre-stim baseline (100%) by one-sample Wilcoxon test: Light OFF Day 1, p=0.063; Light OFF Day 2, p=0.031; Light OFF Day 3, p=0.500. Behavior is normalized to the breakpoint on the day prior to optogenetic inhibition. **E.** Optogenetic inhibition of Gad2 neurons rescued chronic stress effects on reward devaluation behavior four days later (N=5-6 mice/group; **P=0.004 for contrast of Gad2/Cre vs WT controls by Mann Whitney). Behavior is normalized to the breakpoint on the day prior to devaluation testing. **F.** Experimental timeline for 20 Hz optogenetic activation of Gad2 neurons (ChR2 in Gad2/Cre mice). **G.** Activation of Gad2 neurons increased the breakpoint in Gad2/Cre mice (N=6-7 mice/group; *P=0.031, by Wilcoxon rank sum test in Gad2/Cre mice; NS = not significant in wildtype controls). **H.** Similar to (D), this effect persisted for 1 day after the optogenetic activation session before dissipating on Days 2-3 (p=0.010 by Friedman test for repeated measures; *P=0.022–0.046 for Dunn’s post-hoc contrasts of Day 1 vs. Days 2 and 3; NS = not significant for other post-hoc contrasts. # = significantly different from pre-stim baseline (100%) by one-sample Wilcoxon test: Light OFF Day 1, p=0.031; Light OFF Day 2, p=0.906; Light OFF Day 3, p=0.781. Behavior is normalized to the breakpoint on the day prior to optogenetic activation. **I.** Optogenetic activation of Gad2 neurons failed to rescue chronic stress effects on reward devaluation behavior four days later (P=0.524 for contrast of Gad2/Cre vs WT controls by Mann Whitney). Behavior is normalized to the breakpoint on the day prior to devaluation testing. **J.** Experimental timeline for SERT cell inhibition using an inducible anion-conducting channel rhodopsin (IC++) in SERT/flp mice. **K.** Inhibition of SERT neurons had no effect on the breakpoint (N=5-6 mice/group; P=0.125 for SERT-flp mice and P=0.313 for WT mice by Wilcoxon). **L.** Replication of the result in (K) in a second testing session two days later (P=0.500 for SERT/flp mice and P=0.406 for WT mice by Wilcoxon). **M.** Optogenetic inhibition of SERT neurons failed to rescue chronic stress effects on reward devaluation behavior (P=0.849 for contrast of SERT/Cre vs WT controls by Mann Whitney).

Next, we tested whether activating DRN Gad2 cells could increase operating responding in this task. Here, we injected a Cre-dependent channelrhodopsin (ChR2) and implanted an optical fiber in the DRN of Gad2 mice, to specifically activate DRN^Gad2^ neurons during the PR task (**Fig. 4f; Supplementary Fig. 6b,d**), following the same experimental procedure as above. We found that activation of Gad2 neurons increased the breakpoint (**Fig. 4g**), indicating that Gad2 activation is sufficient to drive operant responding in this task. Similar to the optogenetic inhibition experiment, this effect persisted for one day after the optogenetic manipulation before dissipating on Days 2-3 (**Fig. 4h**). As expected, this manipulation did not rescue stress effects on reward devaluation behavior (**Fig. 4i**).

Finally, to determine whether this effect was specific to DRN GABA neurons, we repeated the optogenetic inhibition experiment in serotonergic neurons, expressing the inhibitory opsin iC++ in the SERT neurons in the DRN (**Fig. 4j; Supplementary Fig. 6c**). We found that inhibition of DRN SERT neurons had no effect on the breakpoint in the PR task (**Fig. 4k**)—an effect we replicated by repeating the experiment (**Fig. 4L**)—or on reward devaluation behavior after chronic stress (**Fig. 4m**). In combination with our 2P imaging and transcriptomic profiling data, these results show that chronic stress effects on habit-driven behavior are driven by hyperactivity in GABAergic neurons in the DRN, and that pharmacologic and optogenetic interventions aimed at suppressing activity in DRN Gad2 cells are sufficient to reverse stress effects on habit formation and to drive a stable and persistent switch to goal-directed reward-seeking.

## DISCUSSION

Our study identifies the dorsal raphe nucleus (DRN) as a key regulator of stress-induced shifts in reward-seeking and habit formation. Our findings build on a rapidly growing body of work aimed at identifying neurobiological determinants of stress susceptibility and resilience^57–60^ as well as metabolic and neuroimmune mechanisms influencing hedonic function and depression-related behavior^61–65^. Using a combination of two-photon calcium imaging, single-nucleus RNA sequencing, and optogenetics, we show that DRN neurons encode both effort and reward signals at the single neuron level, suggesting a previously unrecognized role in integrating these task variables within the DRN. Notably, reward responsiveness prior to stress predicted the formation of habit-driven behavior after stress, which was associated with increased reward-related activity. At the molecular level, we identified gene expression changes in DRN GABAergic neurons that facilitate increased GABA release but only in mice that develop habit-driven responding, and implicating stress-induced hyper-activity in DRN GABAergic neurons in the emergence of this behavior. Optogenetic inhibition of these neurons rescued stress effects on habit formation and drove a stable switch in operant responding, providing causal evidence that these neurons contribute to stress-induced behavioral inflexibility.

The dorsal raphe nucleus has been implicated in various aspects of motivational control, including reward-seeking and stress regulation^21,28^. Our results extend this understanding by showing that DRN neurons integrate effort and reward, a computation that may allow for adaptive decision-making under normal conditions, and that chronic stress enhances the co-representation of effort and reward within DRN neurons, indicating stress-induced alterations in these computations. Critically, we find that mice that develop habit-driven behavior after stress have lower reward responsiveness prior to stress and subsequently develop increased reward responses after stress exposure suggesting that DRN activity may predict the formation of stress-induced behavioral inflexibility. This indicates that the DRN, traditionally viewed as a neuromodulatory hub, may play a key role in shaping stress-induced behavioral biases.

Furthermore, our molecular findings reveal that DRN GABAergic neurons undergo stress-induced transcriptional and functional changes that increase GABA release selectively in mice exhibiting compulsive habit-based licking. This aligns with previous reports indicating that stress can induce neurotransmitter plasticity within the DRN, including increased corticotropin-releasing hormone (CRH) receptor expression and neurotransmitter switching in serotonergic neurons toward GABA release^15, 17,37^. Our study implicates plasticity in neurotransmitter release in DRN GABAergic subpopulations and directly links the DRN to stress-induced habit formation. Furthermore, our results indicate that DRN GABA neurons, which have been relatively unexplored in comparison to DRN serotonergic neurons, may play a potent and previously underappreciated role in shaping reward-seeking and habit-driven behavior.

Critically, our optogenetic experiments provide causal evidence that DRN GABA neurons are necessary for stress-induced compulsive behaviors. Inhibiting these neurons in stressed animals restored normal, flexible, effortful reward-seeking behavior, demonstrating that their hyperactivity is a key driver of behavioral inflexibility. Given that DRN serotonergic neurons are known to facilitate behavioral adaptation and flexible decision-making^21,28^, this suggests that stress-induced hyperactivity of DRN GABAergic neurons may drive behavioral inflexibility through local inhibition of DRN serotonin neurons. Alternatively, DRN GABAergic neurons may directly modulate striatal activity, shifting decision-making toward habit-based strategies at the expense of goal-directed control, a hypothesis that is supported by our finding that inhibition of DRN serotonin neurons did not rescue habit-driven behavior.

Our work also identified a host of potential drug targets that could be used to rescue these effects—an approach we validated in a preliminary test involving just one drug as proof-of-principle. Our findings highlight DRN GABA neurons as a promising cellular target for therapeutic interventions aimed at rescuing stress effects on habitual behavior in stress-related neuropsychiatric disorders. Our analysis of receptor expression in these neurons identifies numerous molecular targets that could be engaged by existing small-molecule drugs acting on Oprm1 or other G-protein coupled receptors. Notably, numerous serotonergic interventions have been explored in the treatment of stress-related disorders such as obsessive-compulsive disorder and post-traumatic stress disorder;^38,39^ our results suggest that modulating DRN GABA neurons may offer a novel therapeutic avenue with greater specificity. Together, these experiments reveal a previously unrecognized role for DRN GABAergic neurons in shaping stress-induced habit formation and behavioral inflexibility and provides a foundation for future research into therapeutic strategies targeting these neurons.

## ACKNOWLEDGMENTS

CL was supported by grants from the National Institute of Mental Health, the National Institute on Drug Abuse, the Hope for Depression Research Foundation, the Pritzker Neuropsychiatric Disorders Research Consortium, and the Foundation for OCD Research. CL has served as a scientific advisor to Delix Therapeutics and Brainify.AI. MW was supported by grants from the National Institute of Mental Health, the Brain and Behavior Research Foundation, and the Leon Levy Foundation. All other authors declare no conflicts of interest.

## MATERIALS AND METHODS

### Mice

Male and female mice were aged 9-12 weeks at the beginning of the experiments. Wild type C57BL/6 (Jackson Labs) were used for behavioral experiments, two-photon calcium imaging, and RNA sequencing. Heterozygous SERT-Flp (B6.Cg-Slc6a4tm1.1(flpo)Luo/J) and Gad2-Cre (B6J.Cg-Gad2tm2(cre)Zjh/MwarJ) mice were used for optogenetics manipulations in DRN serotoninergic and GABAergic neurons, respectively. CD1 male mice (Charles River Laboratories) were used as aggressors for chronic social stress experiments. Mice were housed in a Weill Cornell Medicine facility and were maintained on a 12-hour light-dark cycle. Except when water-restricted for behavioral training and testing, all mice were given ad libitum access to food and water. Littermates underwent surgeries within the same week, and mice were group-housed with littermates with the same surgery status. All procedures were approved by the Weill Cornell Medicine IACUC.

### Surgery

Animals were placed inside a flow box and anesthetized with isoflurane gas (3% in 1 l/min oxygen) until sedated, at which point they were placed in a stereotactic device and maintained on 1% isoflurane for the duration of the surgery. Eye gel (Optixcare) was applied to both eyes, and body temperature was maintained at 37 °C with a heating pad and rectal probe. Meloxicam (2mg/kg) and buprenorphine (0.5mg/kg) were injected intraperitoneally. For all surgeries, scalp hair was trimmed away, a midline incision was made using fine surgical scissors (Fine Science Tools) for exposing the skull, and 0.1ml of local anesthetic agent (bupivacaine 0.25%) was topically applied. The skull was then cleaned with 3% hydrogen peroxide and immediately rinsed with saline solution. After surgery, mice received a subcutaneous injection of sterile saline solution and were placed in a recovery cage on a heating pad until they awoke from anesthesia. To avoid post-operative pain, meloxicam was intraperitoneally injected for 3 days.

#### Head fixed surgery

To allow for head fixation for behavioral experiments, a custom-made head post was placed on top of the exposed skull and secured with Metabond (Parkell Inc). Mice were maintained under anesthesia until the Metabond solidified, and the implant was secured.

#### Microendoscopic surgery

Two weeks prior to the microendoscope implantation, 1 µl of AAV9.Syn.GCaMP6s.WPRE.SV40 (Addgene #100843) was injected into the DRN (bregma: AP, -4.5 mm; ML, 0; DV, -3.5 mm). On the day of the microendoscope implantation, a 0.5-mm burr (Fine Science Tools) and a high-speed hand dental drill (Osada) were used to perform a craniotomy around the DRN, taking care not to compress brain tissue or damage the sagittal and transverse venous sinus. After removing the skull, the exposed brain was kept hydrated with sterile saline solution. The dura above the microendoscope implantation site was removed using fine forceps (Fine Science Tools). To facilitate the microendoscope implantation, a needle was inserted above the implantation site (with respect to the bregma: AP, -4.5 mm; ML, ±0.4; DV, -2.8 mm) and slowly removed. Then a custom made microendoscope (9.1 mm ProView^TM^ GRIN lens ending with a 1 mm x 1 mm x 1 mm prism, Inscopix #130-000444) was implanted laterally to the DRN (with respect to the bregma: AP, -4.5 mm; ML, ±0.4; DV, -4 mm) and secured with Vetbond and Metabond (Extended Data2 A). A custom-made head post was placed around the microendoscope to allow head fixation of the mice.

#### Optogenetic surgery

For optogenetics experiments, 1 µl of pAAV-nEF-Coff/Fon-iC++-EYFP (Addgene # 137157), AAV8-nEF-Con/Foff 2.0-iC++-EYFP (Addgene #137156) or AAV8-nEF-Con/Foff 2.0-ChR2-EYFP (Addgene # 137163) was injected in the DRN (bregma: AP, -4.5 mm; ML, 0; DV, -3.5 mm) of SERT-flp or Gad2-Cre mice respectively. A 5mm optical fiber (Thorlabs #CFM12L05-10) was then implanted above the injection site (bregma: AP, -4.5 mm; ML, 0; DV, -3.4 mm) (Extended Data3 A). The fiber implant was secured with Vetbond and Metabond.

### Progressive ratio behavior

Animals recovered 30 days from surgery before being placed on water restriction for four days, after which they typically drank 1mL/day. The behavioral experiments were run in a sound-proof box. During the behavioral task, mice underwent one testing session per day.

#### Habituation to behavioral apparatus

The apparatus consisted of a plastic platform covered with an aluminum tape, on which the mice stand with their heads fixed. A rotative waterspout is located directly in front of the mouse’s mouth for water delivery. The reward spout is equipped with a touch sensor, allowing accurate detection and analysis of lick rates. During three sessions, mice were habituated to drink through the waterspout. The session ended when the mice reached a total of 1,500 licks or 200 trials.

#### Behavior training: fixed ratio

Mice were trained in an operant conditioning task under a fixed ratio-1 schedule of reinforcement (FR) during 30-minute daily sessions. For each trial, the availability of the waterspout was indicated to the animal by a 2 kHz tone played in the sound-proof box. Each active lick (Effort period - no water delivered) triggered the delivery of 12 μl of water across 3 seconds (Reward period). Subsequently, the waterspout was rotated out of the reach of the mice for a 12-second intertrial interval (ITI).

#### Behavioral testing: progressive ratio

Following training, mice were engaged in five consecutive sessions under a progressive-ratio schedule of reinforcement (PR). Under this schedule, the mice were required to progressively increase the number of active licks (effort period) every 4 trials based on the following progression sequence^42,43^ : response ratio (rounded to nearest integer) = (5e^(trial#×^ ^0.2)^) − 5. Thus, progressive ratio reinforcement used the following: 1, 2, 4, 6, 9, 12, 15, 20, 25, 32, 40, 50, 62, 77, 95, 118, 145, 178, 219, etc. The maximal number of active licks performed to reach the final ratio was defined as the breakpoint, a value reflecting animals’ motivation to obtain the reward. 2-photon calcium imaging and optogenetic experiments were conducted during the 30 minutes of the PR task.

#### Devaluation

0.5 ml of water was provided 2-5 minutes before the beginning of the PR task to water-restricted mice to reduce their thirst and test their ability to devalue the reward.

### Two-photon calcium imaging

#### Image acquisition

2-photon calcium imaging^44,45^ was performed via an Olympus 25x 1.00NA objective, with 4mm working distance. All images were acquired using a commercial two-photon laser-scanning microscope (Olympus RS) equipped with a scanning galvanometer and a Spectra-Physics Mai Tai DeepSee laser tuned to 920nm, operating at 120-800mW. Fluorescence was recorded through gallium arsenide phosphide (GaAsP) detectors using the Fluoview acquisition software (Olympus) and a green light emission bandpass filter (Semrock). Calcium signals were acquired at 512x512 pixel resolution with a frame rate of 15Hz. All calcium imaging experiments occurred in awake mice during the 30-min session of progressive ratio task.

#### Image processing

Videos were motion-corrected using custom scripts (Joseph M Stujenske) implemented in MATLAB (Mathworks) to account for motion, which can be more significant in the DRN compared to cortex. After down-sampling (final frame rate 3Hz), videos were denoised using DeepCadRT^32^. Cells were identified and visually inspected using Suite2P ^46^ and neuropil and neuronal calcium traces were extracted. The neuropil signal was scaled and subtracted from the cell signal (FluorescenceCell - (0.7 x FluorescenceNeuropil)). The resulting traces were then z-scored and normalized to the baseline activity on each trial (-5 to -2 seconds from the trial onset). This allowed us to control for any bleaching that occurred during the 30-min session. For analysis, each trial was been divided into “Effort”, “Reward,” or “ITI” task epochs.

### Optogenetics

In the PR task, 4.5-5mW light was delivered to behaving mice via a Plexon 465nm fiber-coupled LED, controlled by Radian software (Plexon). Intermittent light was delivered throughout the 30-min PR task. For DRN^SERT^ and DRN^Gad2^ inhibition, continuous light was delivered on a cycle of 4 sec on and 11 sec off. For DRN^Gad2^ activation, the output parameters were 10 ms, 20 Hz, 4 sec on, and 11 sec off cycle as depicted in Figure 4.

### Chronic stress

#### Chronic social defeat stress

As previously described^47^, male C57BL/6 mice underwent 10 days of chronic social defeat stress (CSDS). Briefly, CD-1 strain mice were housed in rectangular hamster cages divided by a clear perforated Plexiglas divider in the middle. On the first day, intruder C57BL/6J mice were placed in the resident aggressor’s home cage for 5-10 minutes. After a daily defeat session, the intruder was transferred across the perforated divider to the opposite compartment and then was housed in this compartment for the remaining 24 hours. A new aggressor in its own home cage was used in each subsequent daily defeat session to prevent habituation to the resident aggressor. Antibiotic gel (Neosporin) was applied locally to injuries. Following the end of the CSDS, all intruders were single housed in standard cages. Control mice stayed group-housed in standard cages.

#### Coat evaluation

To assess response to chronic stress, we evaluated the coat state of each mouse before and after CSDS. Because the coat state is a function of the frequency and extent of grooming behavior, it is a measure of the animal’s motivation to maintain grooming. The coat state varies from smooth and clear in control animals to bristling with spikes in chronically stressed mice. The mice coat was evaluated based on a previously published protocol^48^ that we adapted for this study. Two experimenters scored the following seven body areas with 0 (well groomed) or 1 (poorly groomed): neck, back, abdomen, tail, genital region, forepaws, and hind paws.

#### Social interaction test

We tested mice in an open-field social interaction assay to analyze social interaction behavior in the presence of a novel CD-1 mouse. Mice were placed in an open field arena for two consecutive 2.5-min trials. In the first trial, the CD-1 was absent, and in the second trial, the CD-1 was present in a perforated Plexiglas container (14 cm x 24 cm) within the social interaction zone as per standard protocols^47^. Behavior was video recorded and EthoVision software (Noldus, Wageningen, Netherlands) was used to measure the time spent in the interaction zone.

#### Elevated-plus-maze

The elevated plus maze (EPM) test consisted of a plus-shaped apparatus with two open and two closed arms (side walls), elevated two feet above the floor. During this five-minute test, the mice were video recorded and EthoVision software (Noldus, Wageningen, Netherlands) was used to quantify the time spent in the open and closed arms.

### Fixed tissue processing and imaging

#### Sliced tissue processing

To verify the location of the microendoscope or the optical fiber, animals were transcardially perfused with ice cold 1xPBS and 4% paraformaldehyde (PFA). The brains were dissected, post-fixed overnight in 4% PFA at 4°C, cryoprotected in 20% sucrose, frozen rapidly, and then stored at -20°C before coronal sectioning with a cryostat (40 mm thick). Samples were mounted on 25x75x1 mm slides (Fisher Superfrost) with DAPI-infused Fluoroshield mounting medium. Images were taken on the Leica DM 5500 B microscope with a Leica DFC360 FX fluorescent camera (7x magnification) and EL600 light source, using a 10x apochromat objective with an NA of 0.4. Images were taken in two channels: a green channel to capture GFP and a blue channel to capture DAPI.

#### Intact brain processing

A sample of brains used in our 2P calcium imaging experiments was cleared and imaged using a lightsheet microscope to generate 3D images of the localization of the transfected cells and microendoscope. To this end, mice were transcardially perfused with 20 mL of ice-cold 1x PBS with 0.2% Heparin Sulfate at a speed of 5 mL/minute. They were then perfused with 20mL of ice-cold 4% PFA in 1xPBS at 5 mL/min. The extracted brains were post-fixed in 4% PFA in 1xPBS for 48 hours at 4°C. Brains were then placed in 20 mL of SHIELD OFF solution (LifeCanvas technologies) for 4 days at 4°C followed by incubation in 20 mL SHIELD ON buffer (LifeCanvas Technologies) for 1 day at 37 °C. Subsequently, brains were placed in 20 mL of Delipidation buffer (LifeCanvas Technologies) for 7 days at 45°C with gentle shaking to remove lipids.

Before imaging, brains were washed in 2x in 1xPBS and then transferred to 50% EasyIndex (LifeCanvas Technologies) in di H20 for 24 hours, then 100% EasyIndex for 24 hours. They were then embedded in 2% EasyIndex gel (2% Ultra Low Melting Point agarose in EasyIndex) according to LifeCanvas Protocols. Brains were imaged on a SmartSPIM axially swept lightsheet microscope using a 3.6x Objective at 2 um z steps. Images were destriped using pystripe (https://github.com/chunglabmit/pystripe), and stitched using Terastitcher (BMC Bioinformatics 2012, 13:316.). Stitched images were downsampled 4x and visualized using ImarisViewer (Andor Technologies) to generate three-dimensional renderings and resampled coronal planes.

### RNA sequencing

#### DRN dissection

Anesthetized mice were decapitated, and the brains were rapidly extracted. 1-mm coronal slices containing the DRN (bregma: AP, -4 to -5 mm) were sectioned using a brain matrix (Zivic Instruments), The DRN was dissected in a dissection buffer containing 1x HBSS, 2.5mM Hepes-KOH pH 7.4, 35 mM Glucose, and 4 mM NaHCO3 using a 1-mm circular punch (Extended Data4 A). Brain samples were then rapidly frozen in liquid nitrogen and stored at -80°C. A total of four animals per group (n=4) each of Control, Goal-Directed, and Habit-Driven DRN were pooled.

#### Single nuclei isolation

Purification of nuclei was achieved as previously described^49,50^, with modifications. Each DRN was homogenized with a glass Dounce homogenizer (Kimble Chase; 1984-10002) in a lysis buffer containing 20 mM HEPES KOH pH 7.4, 150 mM KCl, 5 aaPQmM MgCl2, and the nuclei were pelleted by centrifugation at 1,200xg, 10 min, 4°C. Nuclei were further purified in a 29% iodixanol cushion containing 0.25 M sucrose, 25 mM KCl, 5 mM MgCl2, 20 mM tricine pH 7.8 buffer, and centrifuged at 10,000g for 10 minutes. This step was repeated 2X. Myelin floats were removed during each spin. The resulting nuclear pellet was resuspended in a resuspension buffer (0.25 M sucrose, 25 mM KCl, 5 mM MgCl2, 20 mM tricine pH 7.8, EDTA-free protease inhibitor cocktail (Roche, 11836170001)). Nuclei were counted using the Countess Cell counter (ThermoFisher Scientific). 5000 nuclei per sample was processed using 10X genomics using the Chromium Single Cell 3′ Library & Gel Bead Kit v2 (10X Genomics) as per manufacturer’s protocol. Samples and reagents were prepared and loaded into the chip, and droplets were generated using chromium controllers for droplet generation. Reverse transcription was conducted in the droplets. cDNA was recovered through de-emulsification and bead purification. Pre-amplified cDNA was further subjected to library preparation and libraries were pooled and sequenced on an Illumina Novaseq sequencer, 1.5B Flow Cell, paired-end sequencing, 2 x 50 cycles per flow cell.

#### Single-cell Data processing

The 10X libraries were sequenced on Illumina NovaSeqXplus instrument with paired end reads (28 bp for read 1 and 90 bp for read 2). Sequencing data were processed using the 10X Cell Ranger pipeline v7.1.0. First, cell ranger mkfastq was used to demultiplex the samples and generate FASTQ files. Then, cell ranger count aligned the FASTQ files to the pre-built mouse reference genome (mm10-2020-A) and generated a gene expression UMI counts matrix.

For further analysis, Seurat v4.1.0 was used. Cells were filtered based on several criteria, excluding those with fewer than 200 or more than 2,500 detected genes, fewer than 400 or more than 5,000 detected transcripts, or mitochondrial gene content greater than 5%. Doublets were identified and removed using DoubletFinder v2.0.3, assuming a doublet rate of 0.8% per 1,000 cells.

Normalization of Gene expression UMI counts was performed using a deconvolution strategy with the scran (v1.22.1), scuttle (v1.4.0), and batchelor (v1.10.0) R packages. Cells were pre-clustered with the quickCluster function in scran, and size factors were computed for each cluster using the computeSumFactors function. UMI counts were normalized using these size factors and log-transformed with the logNormCounts function in scuttle. Additional normalization across samples was performed using the multiBatchNorm function in batchelor.

To identify highly variable genes, we applied the FindVariableFeature function in Seurat package v4.1.0, selecting the top 3,000 variable genes while excluding mitochondrial and ribosomal genes. The data were scaled using the ScaleData function, and principal component analysis (PCA) was performed using the RunPCA function. The top 25 principal components (PCs) were selected for downstream clustering analysis.

UMAP dimensionality reduction was carried out using the RunUMAP function in Seurat, with the number of neighbors set to 30 and training epochs set to 500. Cells were clustered into 16 clusters by constructing a shared nearest neighbor (SNN) graph and grouping cells with similar transcriptome profiles using FindNeighbors and FindClusters functions in Seurat, with the resolution parameter set to 0.3. Marker genes for each cluster were identified using the FindMarkers function by comparing cells within and outside of each cluster. Clusters were then assigned based on the identified marker genes of the DRN^33,34^. Low-quality clusters that expressed genes from multiple lineages representing mixed populations such as oligodendrocytes and neurons were excluded from further analysis^33^. This resulted in 15 remaining high-quality clusters.

#### Differential Expression and Pathway Enrichment Analysis

We performed pairwise differential expression analysis among the three following conditions for each cluster using the Wilcoxon rank-sum test implemented in the FindMarkers function from the Seurat R package: Habit-Driven versus Ctrl, Goal-Directed vs Ctrl and Habit-Driven vs Goal-Directed. A minimum of 10 cells per condition per cluster was required to perform the analysis. The differential expression results for each pairwise comparison and cluster were visualized using volcano plots generated with the ggplot2 R package.

We selected differentially expressed genes with a p-value < 0.1 and an absolute average log2 fold change > 0.1 for further analysis. These selected differentially expressed genes were subjected to over-representation analysis against the Reactome database using the enrichPathway functions from the clusterProfiler R package (v4.6.2). The enriched pathways were visualized through bubble plots created with ggplot2.

### Statistical analysis

#### General linear model (GLM) in 2P calcium imaging

For 2P calcium imaging, we used MATLAB’s *fitglm* to identify neurons coding for different features of PR task. We tested several behavioral variables in order to optimize the model. As in previous studies^51^, the behavioral variables used in the final model were those whose removal resulted in a significant degradation of the fit of the model prediction to the data across the population (Supplementary Figure 1F). Improved fits were assessed by comparing the R^2^ for each model with linear mixed-effects models, followed by a paired test across the population of mice. The behavioral variables used in the final model were: *event*: a discrete predictor that at each timepoint indicates the current behavioral event of the trial (effort, reward or ITI); and *lick ratio:* a continuous predictor that at each timepoint takes the value of the number of licks required to complete the trial. We also considered another behavioral variable, *licks:* a predictor that at each timepoint indicate whether a lick was detected. This variable was coded as binary predictor, with a value of ‘1’ in all time points of trials where animals licked or ‘0’ elsewhere. The *licks* variable did not improve the fit and therefore was not included in the final model (Supplementary Figure 1F). GLMs were modeled with normal distributions, as linear functions (no interaction effects), and with the events as intercept. Each individual neuron was tested on a separate GLM.

As previously described^51^, to determine if the contribution of a behavioral variable was statistically significant for each neuron (Figure 1F), we first calculated the F-statistic of the nested model comparison test where the reduced model was the model without that behavioral variable included. We then proceeded to calculate the same statistic on 1000 instances of shuffled data, where shuffling was performed on non-overlapping 3-s bins (to maintain the autocorrelation of the signal). The p-value used for significance was obtained by comparing the value of the original F-statistic to the shuffled distribution, using the Bonferroni correction to account for the number of behavioral variables tested for each neuron; the threshold for significance was a p-value of 0.01 after correction. If a predictor was determined significant after correction for a given neuron, we used the sign of the beta weight to determine whether the neuron was “excited” or “inhibited” by each behavioral variable.

#### Binary decoding with SVM classifiers for 2P calcium imaging

MATLAB’s *fitcsvm* function was used to test the ability to decode, from time series neuronal activity, whether mice were expending effort or receiving a reward at each time point. The models were trained using an SVM decoder with 5-fold cross-validation. On each iteration, data were separated into training (80% of trials) and testing (20% of trials) sets, balanced for task epoch. Decoding accuracy was expressed as the percentage of correct predictions per mouse. To generate a null distribution of model performance in shuffled data, we tested the ability of the model to decode the same behavioral events after shuffling (1,000 iterations) the neuronal activity in non-overlapping 3-s bins (to maintain the autocorrelation of the signal) using *randswap*.

#### Linear decoding with SVM classifiers for 2P calcium imaging

MATLAB’s *fitrlinear* function was used to test the ability to decode, from the peak neuronal activity of DRN neurons, the distance to the breakpoint (-log(breakpoint distance)) on any given trial within a testing session. The linear regression models were trained using with 5-fold cross-validation regularized using LASSO. On each iteration, data were separated (bootstrapped) into training (80% of trials) and testing (20% of trials) sets. Decoding accuracy was expressed as the correlation between predicted breakpoint distances and real values. In this way, we trained decoders to predict the breakpoint based on the peak activity during the effort, reward or ITI periods. Performance in shuffled data was estimated as described above. The breakpoint distance was modeled as follows:

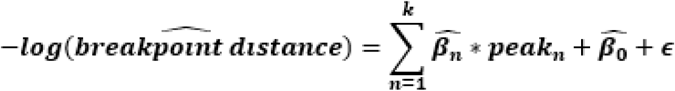

#### Other statistical tests

All statistical analyses were performed using Matlab and GraphPad Prism using non-parametric tests suitable for relatively smaller samples that may not be normally distributed and sample sizes similar to those reported in previous publications^52^. The statistical tests used in each figure panel are reported in the corresponding figure legends (e.g. Wilcoxon, Mann-Whitney, Friedman, Kolmogorov-Smirnov, or linear mixed effects analysis). For linear mixed effects models, mouse or neuron identity was modeled as a random effect. Bonferroni corrections were performed for multiple comparisons.

## Data Availability

All data necessary to understand and evaluate the conclusions of this paper are provided in the Article and Supplementary Information. All RNA-seq data will be deposited upon acceptance of the manuscript.

## Code Availability

Preprocessing of two-photon calcium imaging data used the Suite2P package, which is publicly available at https://github.com/MouseLand/suite2p. Other analyses used standard methods and functions readily available in Matlab (versions 2022b and 2023a).

## Supplementary Material

**Supplementary Figure 1.**
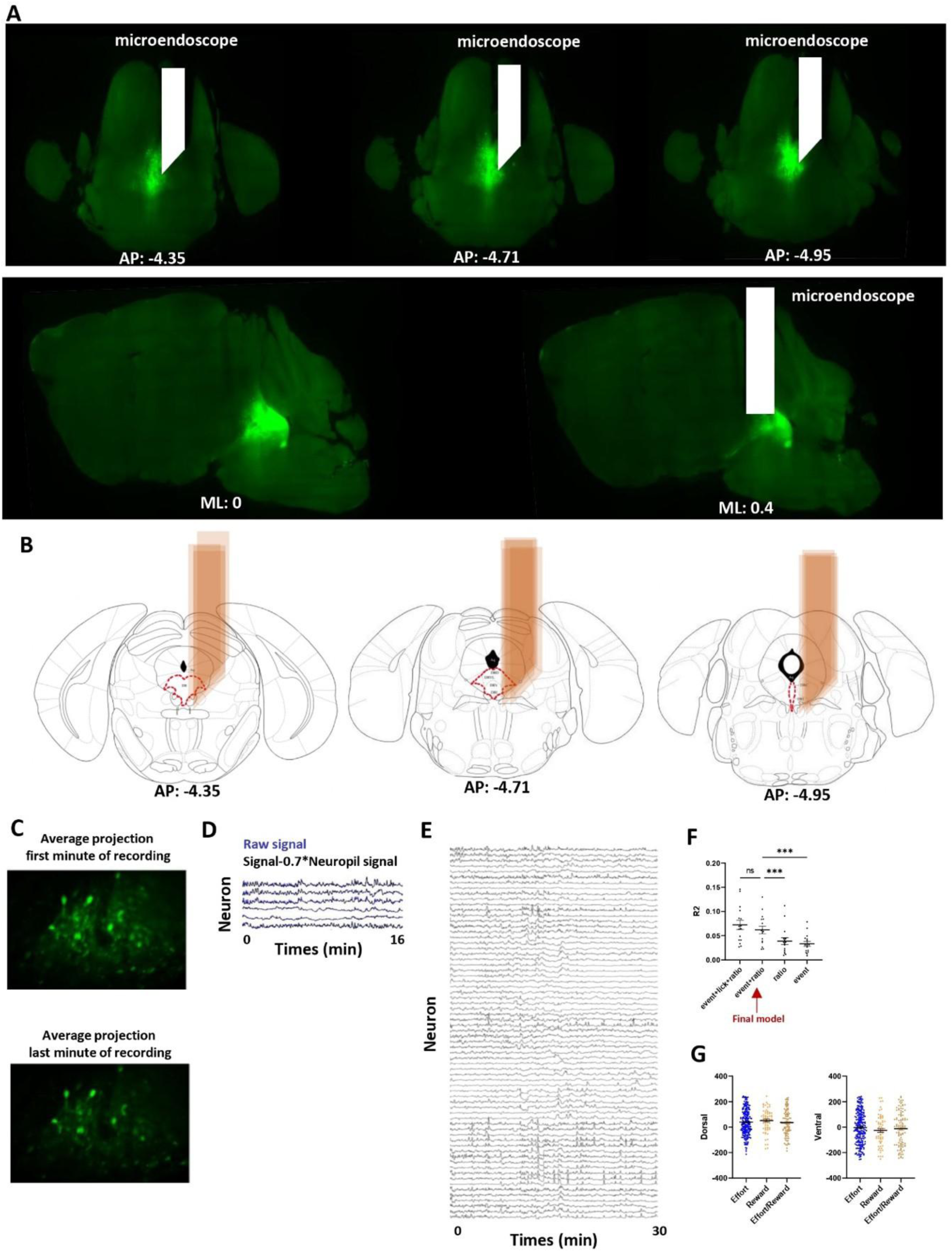
Two-Photon Imaging of DRN Neurons During Progressive Ratio Task Performance. **A.** Example of virus and microendoscope localization for 2P calcium imaging across three coronal views (**top**) and two sagittal views (**bottom**). **B.** Schematic of microendoscope placement of 2P calcium imaging mice. **C.** Mean projection image over the first (**top**) and last (**bottom**) minute of a representative 30-minute recording session. There was no substantial shift in the field of view over time. **D.** Preprocessing of representative calcium signal traces. Raw signal in blue. Signal corrected by neuropil signal in black. **E.** Representative traces from multiple neurons during a single 30-minute recording session. **F.** Variance explained by the GLM model used in our analyses (averaged across animals) compared to other models in which a variable was either removed or added. See Materials and Methods for descriptions of all variables that were tested. All models for which a variable was removed from the final model performed significantly worse in terms of variance explained (R^2^). For models in which a variable was added to those in the final model, the performance did not exhibit a significant difference (n=16 mice, Mixed-effects analysis: Model effect F(6,90)=22.87 p<0.001, Bonferroni corrected post-hoc for model comparisons to the final model: Event p<0.001, Ratio p<0.001, Event+Lick+Ratio p=0.250). **G.** Cells coding for effort, reward and ITI did not differ with respect to their position along the dorso-ventral (**left**) or antero-posterior (**right**) axes of the DRN (n=225 effort cells, n=65 reward cells, n=104 multimodal effort/reward cells; Kruskal-Wallis test for position along dorsoventral axis: F(2,391)=2.077 p=0.354; Kruskal-Wallis test for position along antero-posterior axis: F(2,391)=1.640 p=0.441). Swarm plots represent mean (line) ±SEM and individual datapoints (dots).

**Supplementary Figure 2.**
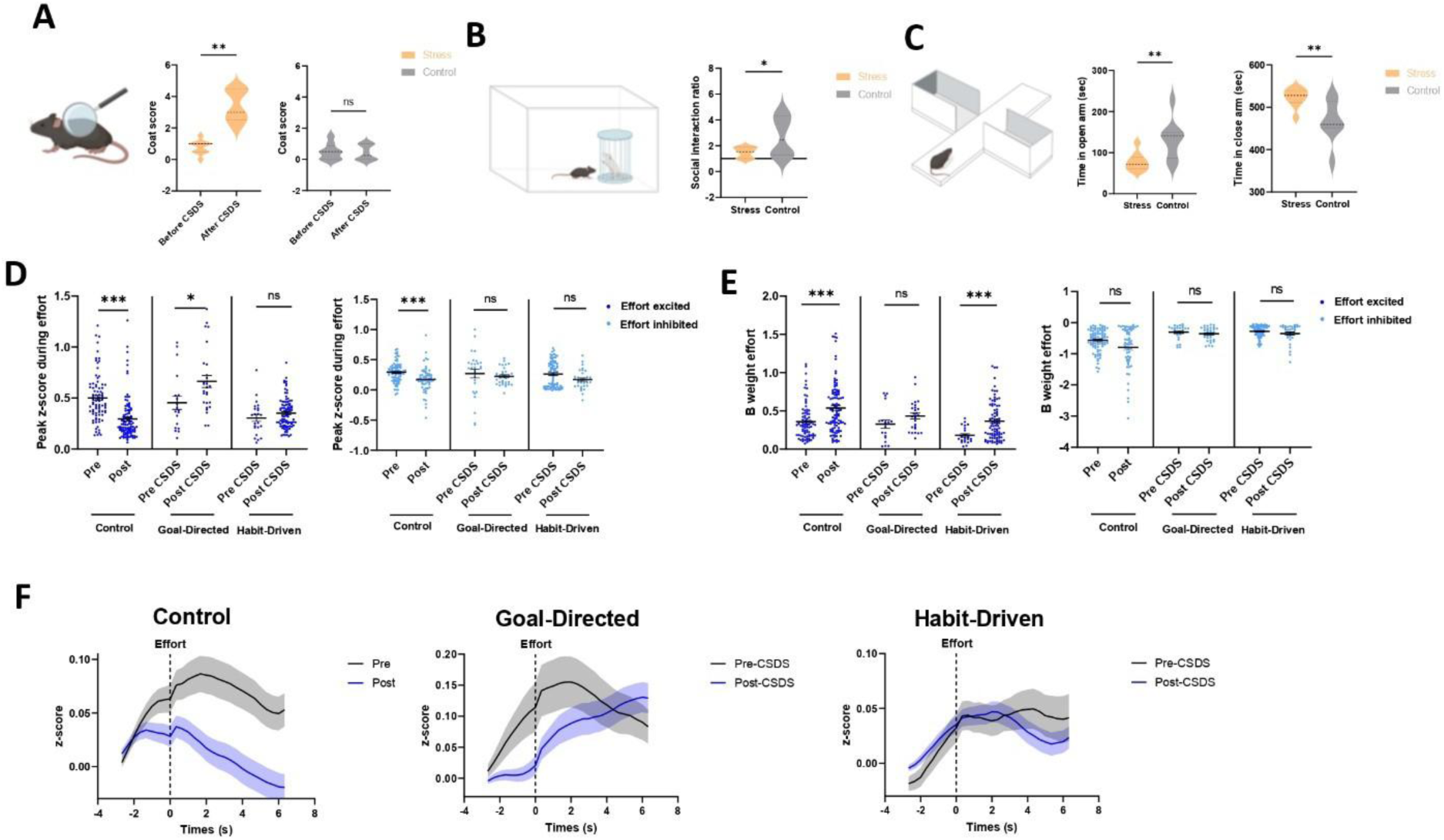
Supplemental Data on Chronic Stress Effects on Behavior and DRN Function. **A.** Effect of chronic social defeat stress (CSDS) on coat evaluation. **Left:** Schematic. **Right:** Coat score increased in stressed mice (n=9, Wilcoxon Test W=45 p=0.004) but not in control mice (n=9, Wilcoxon Test W=-2 p>0.999). **B.** Effect of CSDS on social interaction test behavior. **Left:** Schematic. **Right:** Stressed mice (n= 8) spent less time in the interaction zone with the CD-1, compared to control mice (n=6; unpaired t test t=2.180 p=0.049). **C.** Effect of CSDS on elevated plus maze behavior. **Left:** Schematic. **Right:** Stressed mice (n=8) spent less time in the open arm (Mann-Whitney W=9 p=0.008) and more time in the closed arm (Mann-Whitney W=9 p=0.008) than control mice (n=6). **D.** Effect of CSDS on coding properties of effort coding neurons. In effort-excited neurons, the peak activity during effort expenditure decreased after the handling control procedure in unstressed controls (pre n=84 cells and post n=105 cells in 6 mice), increased after CSDS in stress-resilient / goal-directed mice (pre-CSDS n=19 cells and post-CSDS n=27 cells in 3 mice) and did not change after CSDS in stress-susceptible habit-driven mice (pre-CSDS n=23 cells and post-CSDS n=91 cells in 3 mice). Bonferroni corrected Mann-Whitney Test for pre vs post: control mice: W=1801, p<0.001; goal-directed mice: W=136, p=0.020, habit-driven mice: W=846 p=0.477. In effort-inhibited neurons, the peak activity during effort expenditure decreased after the handling control procedure in unstressed controls (pre n=86 cells and post n=60 cells in 6 mice) and did not change after CSDS in either goal-directed (pre-CSDS n=28 cells and post-CSDS n=34 cells in 3 mice) or habit-driven mice (pre-CSDS n=84 cells and post-CSDS n=34 cells in 3 mice). Bonferroni corrected Mann-Whitney test for pre vs post: control mice: W=1535, p<0.001; goal-directed mice: W=362, p=0.326; habit-driven mice: W=1172, p=0.388. **E.** In GLM analysis, the beta weight for the effort predictor for effort-excited neurons increased after the handling control procedure in unstressed controls and after CSDS in habit-driven mice but not in goal-directed mice. Bonferroni corrected Mann-Whitney test for pre vs post: control mice: W=2856, p<0.001; goal-directed mice: W=179, p=0.258; habit-driven mice: W=544, p<0.001. No difference was observed in effort-inhibited neurons. Bonferroni corrected Mann-Whitney Test for pre vs post: control mice: W=2242, p=0.540; goal-directed mice: W=394, p=0.753; habit-driven mice: W=1302, p>0.999. **F.** Mean activity traces (+/- SEM) time-locked to effort onset, and averaged across all trials and all neurons from (G-H) with significantly positive effort-associated beta weights in unstressed controls (**left**), stress-resilient / goal-directed mice (**middle**), and stress-susceptible / habit-driven mice (**right**). Swarm plots represent mean (line) ±SEM and individual datapoints (dots). Violin plots represent mean (dashed lines), first and third quartiles (dotted lines). Line plots represent mean (line) ±SEM (shaded area).

**Supplementary Figure 3.**
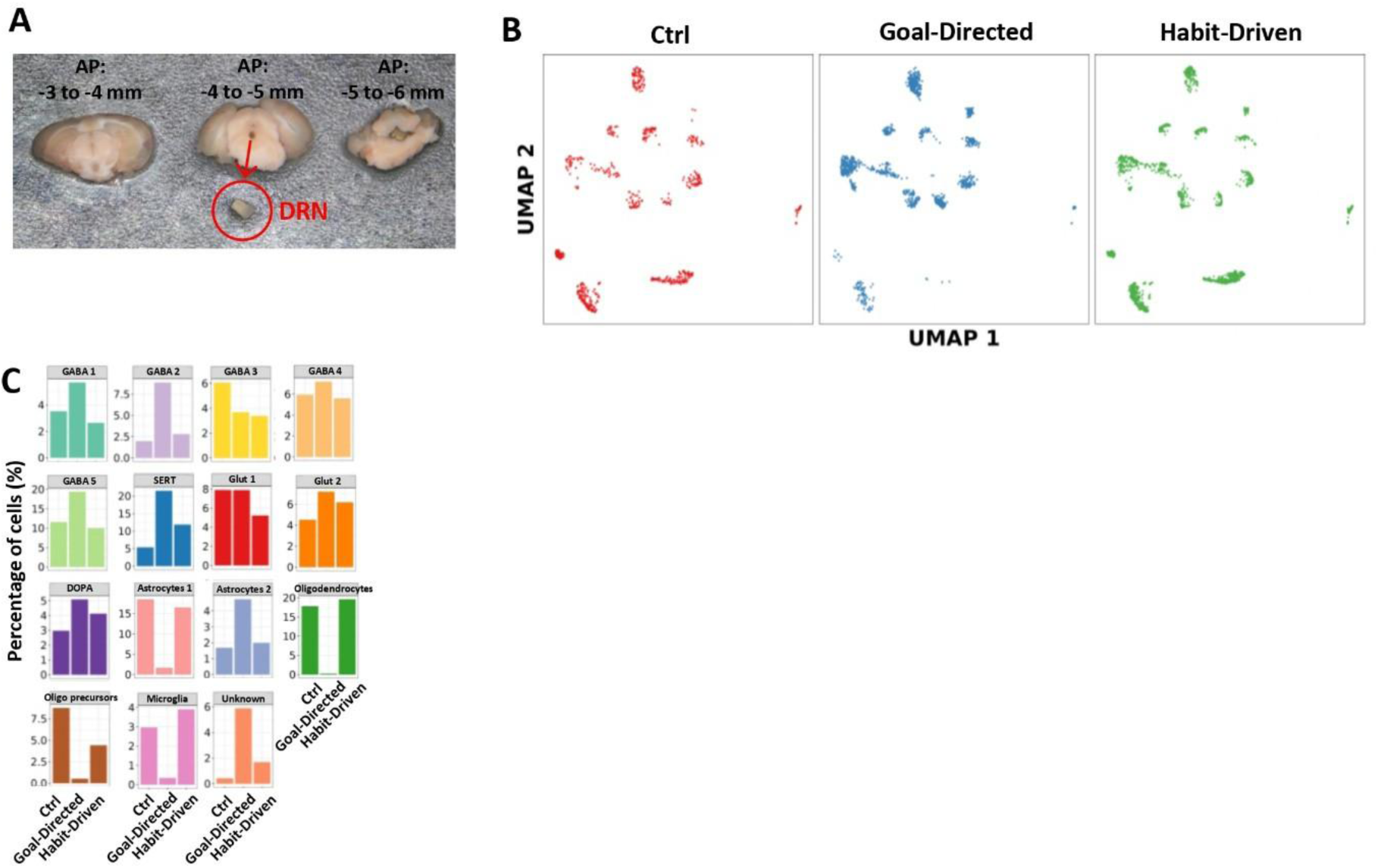
Supplementary Data on Transcriptomic Profiling Experiment. **A.** Representative DRN dissection using 1mm punch. **B.** UMAP visualization of the single-nuclei clusters across the three experimental groups (Control, Goal-Directed, Habit-Driven). **C.** Histograms of percentage of cells constituting each cluster (n=15 clusters) across experimental groups (Ctrl, Goal-Directed, Habit-Driven). There was no difference between experimental groups (Friedman Test, F(2,42)=2.533 p=0.282).

**Supplementary Figure 4.**
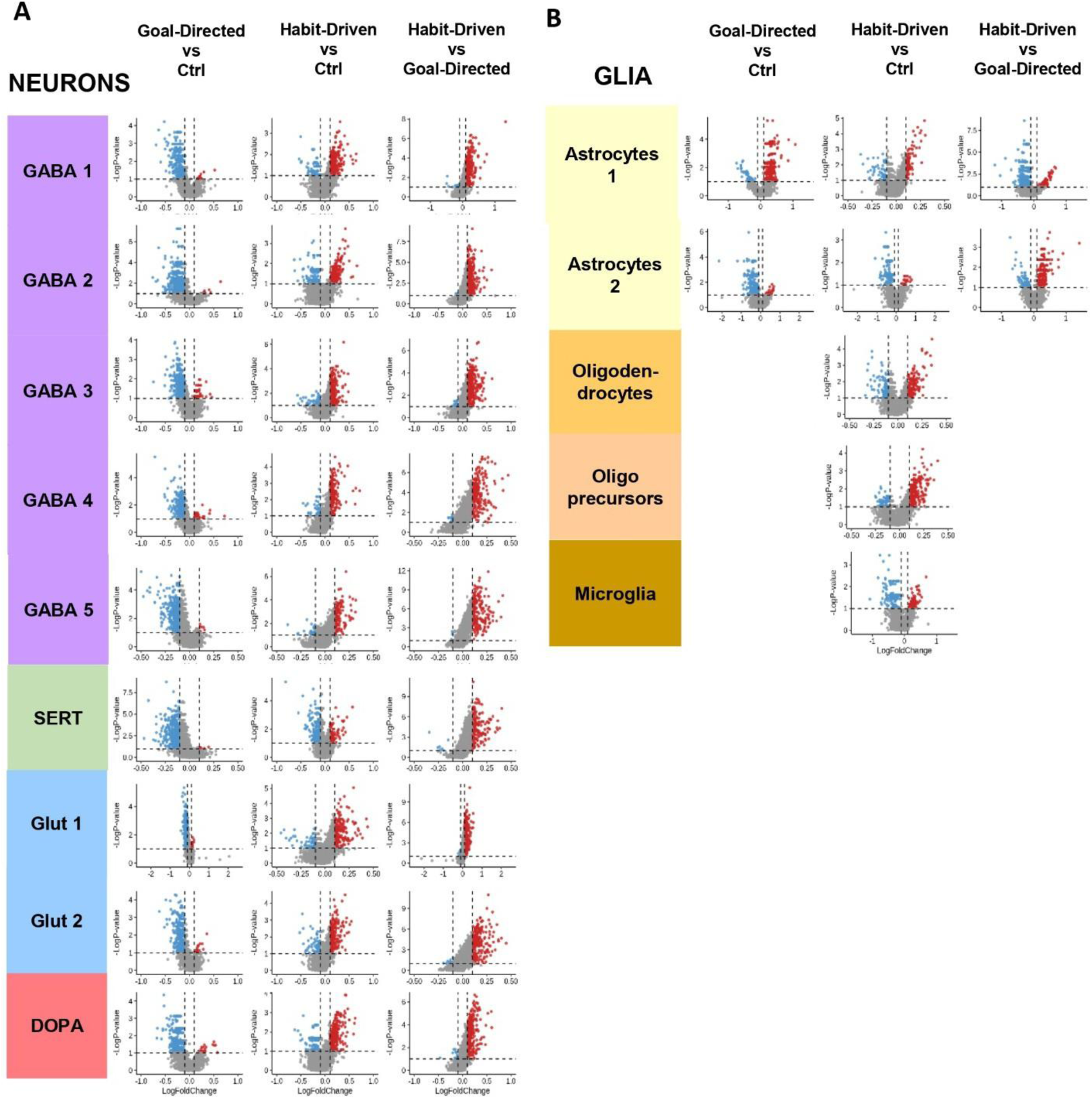
Supplementary Differential Gene Expression Data. Volcano plot illustrating differential gene expression between three experimental groups in neuronal (**A**) and glial (**B**) clusters in stress-susceptible / habit-driven mice, stress-resilient / goal-directed mice, and unstressed control mice (Ctrl). Upregulated genes are represented in red, downregulated genes in blue, and genes without significant changes in gray.

**Supplementary Figure 5.**
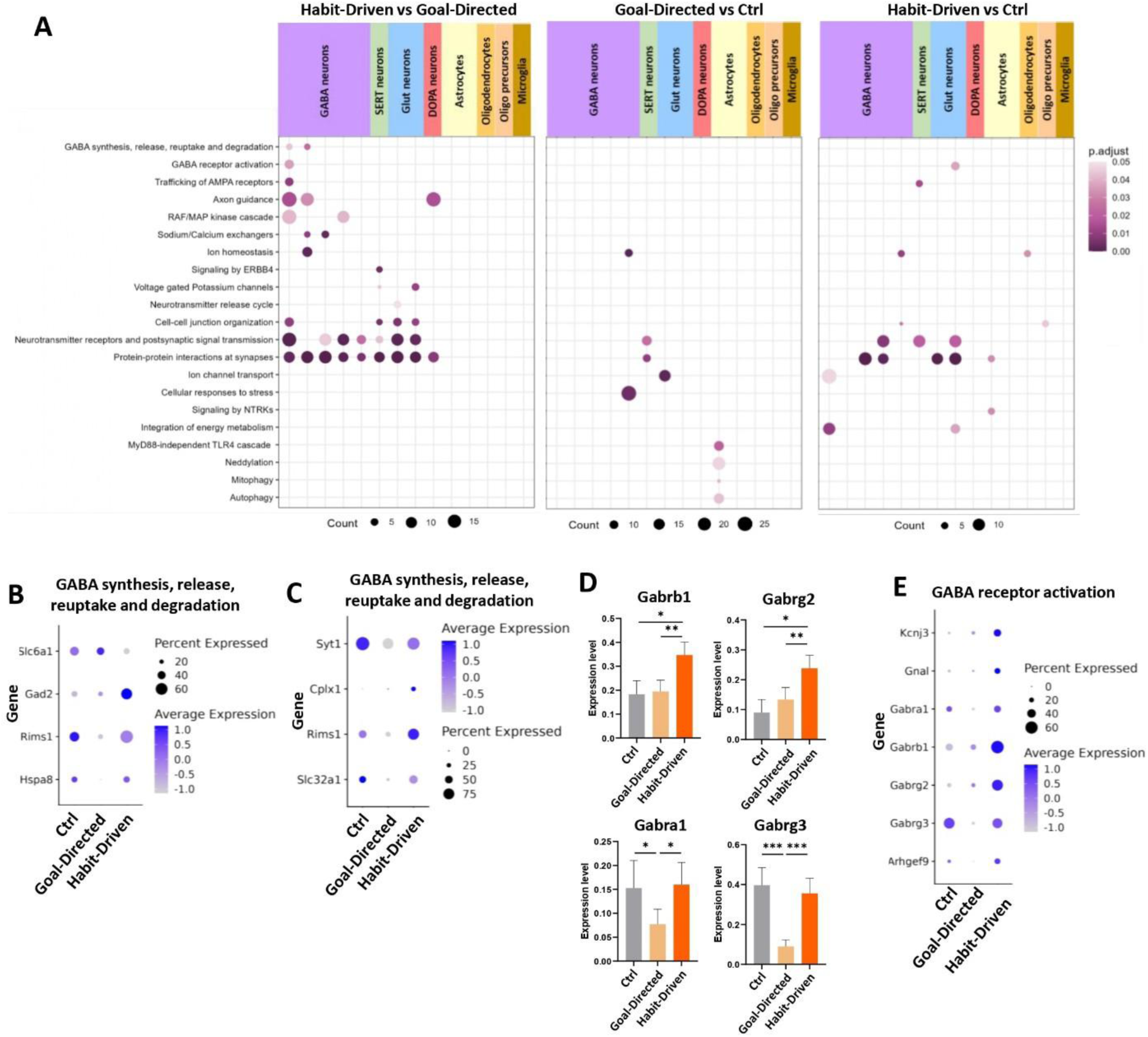
Supplementary Data on Pathway Analysis. **A.** Top enriched pathways of differentially expressed genes identified in Habit-Driven vs Goal-Directed mice (**left**), Goal-Directed vs Control mice (**middle**) and Habit-Driven vs Control mice (**right**). The color gradient of each dot indicates the adjusted p-value of each pathway, whereas the dot size indicates the number of significant gene in the pathway. **B.** Dot plots of expression of differentially expressed genes of “GABA synthesis, release, reuptake and degradation” pathway in GABA neuron cluster 1. The color gradient of each dot indicates the average expression of each gene whereas the dot size indicates the fraction of cells expressing the gene. **C.** Dot plots of expression of differentially expressed genes of “GABA synthesis, release, reuptake and degradation” pathway in GABA neuron cluster 2. The color gradient of each dot indicates the average expression of each gene whereas the dot size indicates the fraction of cells expressing the gene. **D.** Bar plots representing the expression level of GABA receptor subunits (*Gabrb1, Gabrg2, Gabrg3, Gabra1*) in GABA neuron cluster 1, compared across experimental groups. The expression level of *Gabrb1* (Wilcoxon test: Goal-Directed vs Ctrl p=0.0.689, Habit-Driven vs Ctrl p=0.043, Habit-Driven vs Goal-Directed p=0.005) and *Gabrg2* (Wilcoxon test: Goal-Directed vs Ctrl p=0.847, Habit-Driven vs Ctrl p=0.013, Habit-Driven vs Goal-Directed p=0.002) was higher in the Habit-Driven group in comparison to the Goal-Directed and Ctrl groups. The expression level of *Gabrg3* (Wilcoxon test: Goal-Directed vs Ctrl p<0.001, Habit-Driven vs Ctrl p=0.688, Habit-Driven vs Goal-Directed p<0.001) and *Gabra1* (Wilcoxon test: Goal-Directed vs Ctrl p=0.086, Habit-Driven vs Ctrl p=0.738, Habit-Driven vs Goal-Directed p=0.013) was lower in the Goal-Directed group in comparison to the Habit-Driven and Ctrl groups. **E.** Dot plots of expression of differentially expressed genes of “GABA receptor activation” pathway. The color gradient of each dot indicates the average expression of each gene whereas the dot size indicates the fraction of cells expressing the gene.

**Supplementary Figure 6.**
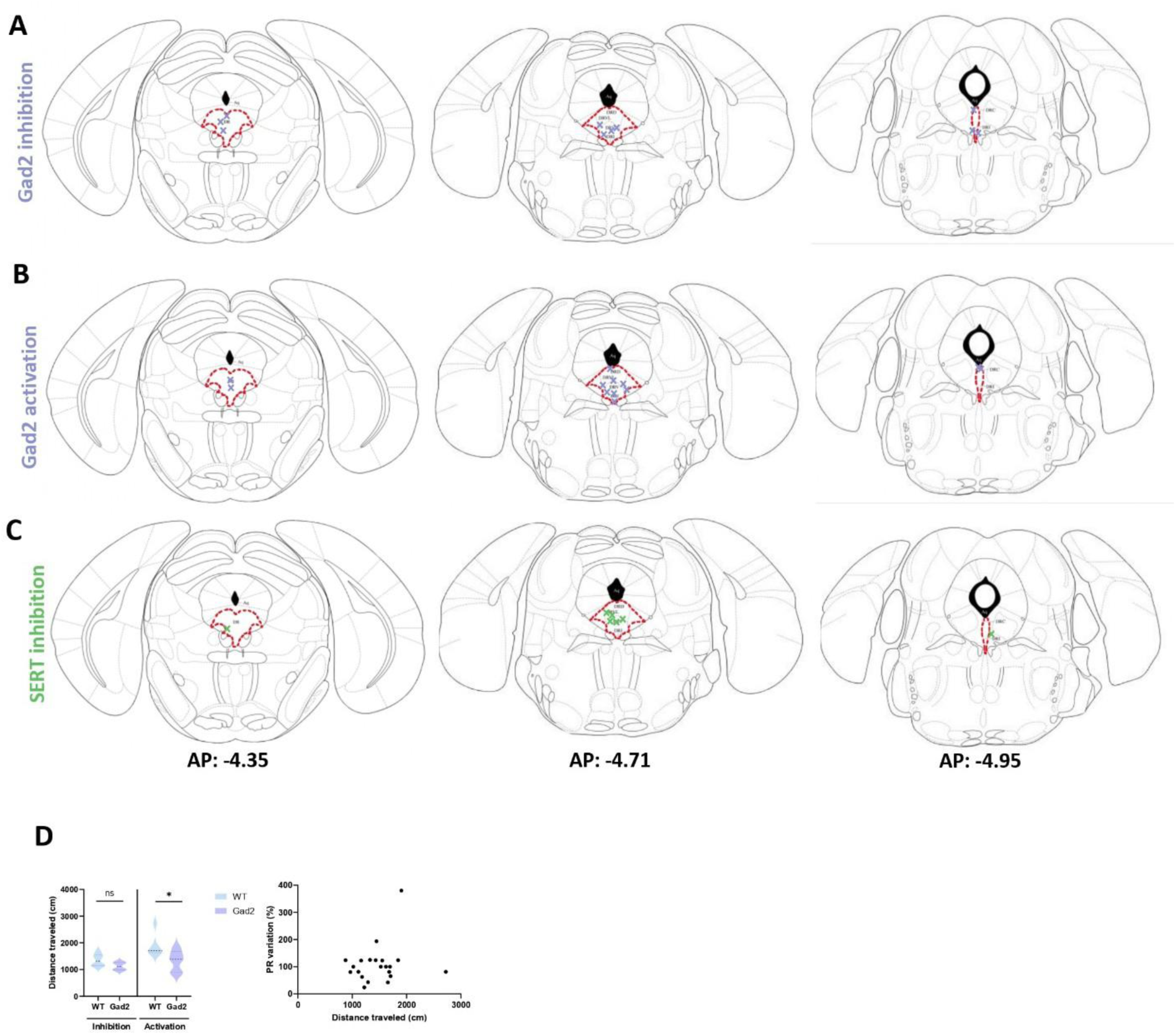
Supplementary Data on Optogenetic Experiments in Fig. 4. **A.** Optical fiber placement for optogenetic inhibition of Gad2 neurons. **B.** Optical fiber placement for optogenetic activation of Gad2 neurons. **C.** Optical fiber placement for optogenetic activation of SERT neurons. **D. Left:** Effect of the optogenetic manipulation in Gad2 neurons on locomotor activity. Optogenetic activation (n=6 WT mice, n=6 Gad2 mice), but not optogenetic inhibition (n=6 WT mice, n=4 Gad2/Cre mice) of Gad2 neurons reduced locomotor activity as measured by a reduced distance traveled in open field (2-way ANOVA: main effect of optogenetic manipulation: F(1,18)=6.218, p=0.023; main effect of genotype: F(1,18)=6.961, p=0.017; optogenetic manipulation x genotype interaction: F(1,18)=1.166, p=0.295. Bonferroni corrected post-hoc contrasts for WT vs Gad2/Cre: inhibition p=0.619, activation p=0.024. **Right:** There was no correlation between distance traveled and behavior in the PR task induced by the optogenetic manipulation (linear regression R^2^=0.028, p=0.454), indicating that non-specific effects on locomotor activity were unlikely to have influenced the effects of optogenetic manipulations on PR behavior. Violin plots represent mean (dashed lines), first and third quartiles (dotted lines). Scatter plots represent individual datapoints (dots).

